# LIPL-1 and LIPL-2 are TCER-1-regulated Lysosomal Lipases with Distinct Roles in Immunity and Fertility

**DOI:** 10.1101/2025.07.14.664648

**Authors:** Laura Bahr, Francis RG Amrit, Paige Emily Silvia, Danny Bui, Bella Wayhs, Mirae Choe, Guled Osman, Nikki Naim, Margaret Champion, Jiali Shen, Javier E Irazoqui, Carissa Perez Olsen, Arjumand Ghazi

## Abstract

Reproduction and immunity are fundamental, energy intensive processes that often compete for resources, leading to trade-offs observed across diverse species. Lipid metabolism plays a crucial role in integrating these processes, particularly during stressful conditions such as pathogenic infections. Yet the molecular mechanisms governing this integration remain poorly understood. TCER-1, the *C. elegans* homolog of mammalian TCERG1, suppresses immunity and promotes fertility, especially upon maternal infection. Here, we show that TCER-1 regulates two conserved lysosomal lipases, *lipl-1* and *lipl-2*, to balance reproduction, immunity and lifespan. Using transcriptomic, lipidomic, and molecular-genetic analyses, we demonstrate that while both *lipl-1* and *lipl-2* mediate infection-induced lipid remodeling, *lipl-1* enhances immunity and catalyzes the accumulation of ceramide species linked to stress response and longevity, whereas, *lipl-2* unexpectedly does not. Both lipases contribute towards fertility outcomes, but *lipl-2* is especially critical for maintaining embryonic-eggshell integrity during maternal infection and aging. Strikingly, expression of human lysosomal acid lipase (LAL), the ortholog of *lipl* genes, rescues the immune defects triggered by *lipl-l* loss and enhances immune resilience. Together, these findings uncover functionally distinct roles for *lipl-1* and *lipl-2* in modulating lipid species that shape immune fitness, healthspan and reproductive health, and suggest a potentially conserved mechanism by which lipid metabolism links fertility and immunity.

## INTRODUCTION

Reproduction and immunity are intimately interconnected across species, from insects to humans, and often exhibit antagonistic interactions. In many species, infections reduce fertility, whereas, increased reproductive effort suppresses immune fitness [1, 2]. However, exceptions exist as mating in some species triggers beneficial immune remodeling and heightened immune response [3, 4], and male-derived factors can confer a survival advantage on females during post-mating infections [5]. In women, pregnancy was once considered to be a state of generalized immunosuppression but is now recognized to involve finely tuned immune adaptations that are essential from implantation through parturition, yet may also impair defense against certain pathogens [6–9]. Importantly, the intense metabolic demands of immunity and reproduction necessitate that these processes are tightly coregulated with energy metabolism [10, 11]. Indeed, immune remodeling during pregnancy is partially driven by lipid metabolism, whereas, dysregulated energy metabolism underlies many pregnancy complications, including gestational diabetes and preeclampsia, the most common pathologies of human pregnancy in the developed world [12–14]. Thus, lipid metabolism emerges as a crucial arbiter of reproductive fitness and immune health coordination.

The molecular mechanisms linking lipid remodeling, immunity, and reproduction are poorly understood. In mammals, complexities of pregnancy, low fecundity, and long gestations hinder mechanistic studies, whereas, they have been more tractable to evolutionary investigations in invertebrates [1, 15]. The nematode model, *Caenorhabditis elegans,* offers unique advantages because its short generation time and high reproductive output (∼300 eggs over 3–6 days) allow sensitive detection of even minor fertility shifts under immune stress and *vice versa* [15–18]. For instance, maternal bacterial infection rapidly mobilizes lipids to the germline to support progeny but results in diminished maternal immune resistance [19–21]. Moreover, despite a simple immune system, *C. elegans* retains conserved immune signaling pathways, and many established infection models exist [22–27].

Previously, we identified TCER-1, *C. elegans* homolog of a mammalian transcription elongation and splicing factor, TCERG-1 [28, 29], as a longevity-enhancing protein, that promotes fertility while suppressing immune responses in reproductively active animals [16]. Pertinently, TCER-1 supports reproductive success, especially during immune challenges, as *tcer-1* loss-of-function (*lof*) mutants exhibit reproductive defects, whereas, TCER-1 overexpression partially mitigates the fertility loss associated with maternal infection [16]. TCER-1’s role in coordinating reproduction, immunity, and aging appears to be evolutionarily conserved as inactivation of its *Arabidopsis* homologue, PRP40, enhances tolerance towards pathogens and delays flowering [30]. Mammalian *TCERG1* is highly expressed in reproductive tissues, and similar to *C. elegans* TCER-1, declines with age in mouse and human oocytes [31, 32]. We showed that TCER-1, along with pro-longevity transcription factors, DAF-16 and NHR-49, increases lifespan upon germline loss through concomitant enhancement of lipid anabolism and catabolism [33, 34]. We have now discovered that TCER-1 impairs immunity and promotes fertility, in part through regulation of lysosomal lipases, *lipl-1* and *lipl-2*-two of eight members of the conserved *‘lipl’* lipase family orthologous to the lysosomal acid lipase (LAL) in humans [35, 36]. The *‘lipl’* genes have been linked to nutrient stress responses, with *lipl-1* and *lipl-3* reported to mobilize fats during starvation [37], *lipl-2* and *lipl-5* to mediate metabolic remodeling upon dietary restriction [38–40] and *lipl-4* to promote longevity by activating a lysosomal-nuclear retrograde signaling pathway [41–43]. Notably, *lipl-1*, *lipl-2*, *lipl-3*, and *lipl-5* have been reported to be induced by pathogens such as *Enterococcus faecalis, Staphylococcus aureus, Cryptococcus neoformans,* and *lipl-2* is differentially expressed during *Bacillus thuringiensis* infection [43–45]. Yet, their functional roles in innate immune response remain unknown.

In this study, we show that TCER-1 regulates *lipl-1* and *lipl-2* expression upon infection by the human opportunistic Gram-negative pathogen, *Pseudomonas aeruginosa* strain PA14 (PA14) [27, 46]. These lipases perform distinct functions in influencing immunity, reproduction, and longevity, and exert discrete effects on the animal’s lipid composition. *lipl-1* enhances pathogen resistance, whereas *lipl-2* does not and, in fact, appears to limit immunity and longevity. Both lipases contribute towards fertility success, but *lipl-2* is especially critical for maintaining embryonic-eggshell integrity during maternal infection and aging. Notably, LIPL-1 and LIPL-2 influence the levels of individual lipid species that have been linked to stress resistance, lifespan, healthspan and reproductive outcomes. Lastly, we show that human LAL rescues *lipl-1 lof*-associated immune defects and enhances worm survival upon PA14 infection. Altogether, these findings provide evidence for discrete roles of LIPL-1 and LIPL-2 in *C. elegans* fertility and immunity, and as central effectors of TCER-1 in coordinating immune, reproductive, and aging processes.

## RESULTS

### TCER-1 broadly remodels lipid metabolism in response to pathogenic infection in *C. elegans*

To elucidate the transcriptomes dictated by TCER-1 under normal conditions and upon pathogenic infection, we performed RNA-Seq on wild-type (WT) animals and *tcer-1(tm1452)* mutants that were exposed to PA14 for 8h and those maintained on standard *Escherichia coli* strain OP50 (OP50) lawns (Fig. 1A). Analysis of the sequencing data using the CLC Genomics Workbench (Table S1; see Methods for details) revealed that pathogen exposure triggered differential expression of 1002 genes in WT animals (Table S1A) of which 583 were upregulated (Table S1Ai) and 419 were downregulated (Table S1Aii). *tcer-1* mutants showed differential expression of 159 genes as compared to WT on the normal OP50 diet (Table S1B) with 81 being upregulated (Table S1Bi) and 78 being downregulated (Table S1Bii). The expression of 306 genes was altered upon comparing WT animals and *tcer-1* mutants after PA14 exposure, constituting a newly identified group of TCER-1-dependent, PA14-responsive genes (Table S1C), of which 150 were upregulated (Table S1Ci) and 156 were downregulated (Table S1Cii). 962 genes were differentially expressed by *tcer-1* mutants on OP50 *vs.* PA14 (Table S1D) with 707 upregulated (Table S1Di) and 255 downregulated (Table S1Dii) and classified as TCER-1-independent, PA14-responsive genes (Fig. 1B).

**Fig 1:**
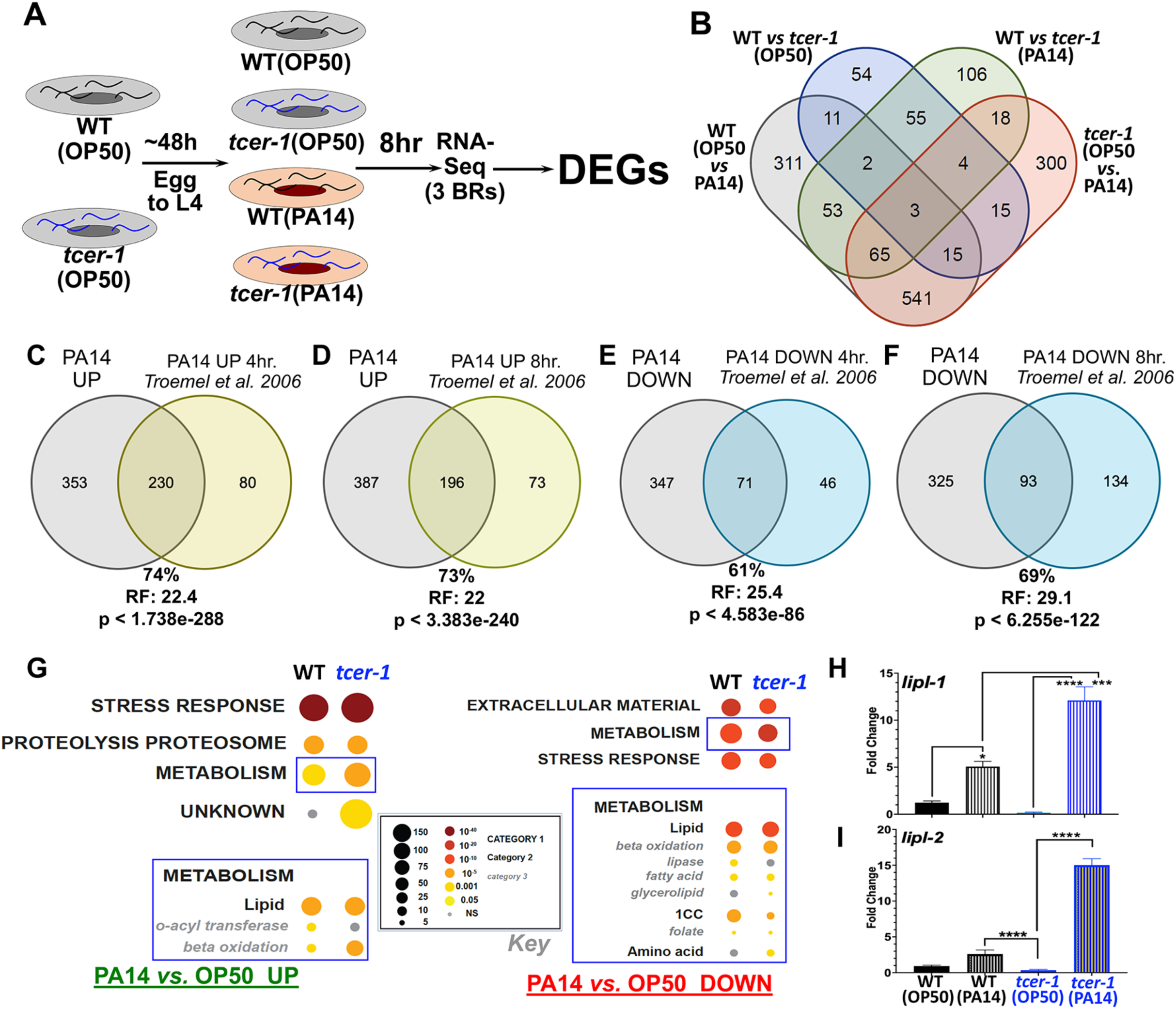
TCER-1 regulates lipid metabolism upon PA14 infection. **A) RNAseq experimental paradigm.** Age-matched wild type (WT) and *tcer-1* L4s raised on OP50 exposed to PA14 for 8 hours. **B) Overlaps of groups of differentially expressed genes (DEGs) identified. C-F)** Overlap of genes upregulated **(C, D)** and downregulated **(E, F)** upon PA14 exposure with previously-identified PA14-responsive genes by Troemel et al. 2006. RF: Representation Factor. Statistical significance of overlap between gene sets calculated using hypergeometric probability formula with normal approximation (see Methods). Comparisons with additional studies in Fig. S1 and Table S2. **G) Gene ontology (GO) term analysis of PA14- and TCER-1-driven DEGs using WormCat.** Metabolism (Category 1), particularly lipid metabolism, (Category 2) (blue boxes) identified one of the most differentially impacted processes amongst genes upregulated (UP, left) and downregulated (DOWN, right) in WT and *tcer-1* mutants. Key shown in middle (black box). **H, I) *lipl-1* and *lipl-2* are transcriptionally upregulated in *tcer-1* mutants on PA14 infection.** mRNA levels of *lipl-1* **(H)** and *lipl-2* **(I)** measured by qPCR in WT (black) and *tcer-1* mutant (blue) adults maintained on OP50 (solid bars) or exposed as L4s to PA14 for 8 h (hashed bars). Data from at least 3 independent trials/biological replicates. Asterisks represent statistical significance of differences observed in unpaired, two-tailed t-tests with P values * ≤ 0.05, *** p≤0.001 and ****p≤0.0001.

PA14-induced gene expression changes have been described in previous reports, and we found a substantial overlap between these datasets and our PA14-induced gene lists. For instance, 230 of 310 genes (74%) found to be upregulated 4 hours after PA14 infection by Troemel et al., were also upregulated in our study (Representation Factor (RF): 22.4, p < 1.738e-288) as were 196 of 269 genes (73%) upregulated after 8 hours of infection (RF: 22.0, p < 3.383e-240) (Fig. 1C, D, Table S2) [27]. 71 of 117 (61%; RF: 25.4, p < 4.583e-86) and 93 of 134 (69%; RF: 29.1, p < 6.255e-122) genes downregulated post-infection at 4 hours and 8 hours, respectively, in the same study were also downregulated on PA14 in our experiment (Fig. 1E, F, Table S2) [27]. Similarly significant overlaps were observed upon comparison with PA14-dependent genes identified by other studies (Fig. S1, Table S2) [26, 47]. These comparisons reinforced our confidence in the transcriptomes that we mapped, including the hundreds of newly-identified PA14 responsive genes, as well as the TCER-1 downstream targets.

Gene Ontology (GO) analysis revealed the PA14-and TCER-1-dependent DEGs to be highly enriched for innate immunity, stress response and metabolism, especially lipid-metabolic functions, as previously noted (Table S1A-D) [33, 44]. Using WormCat, a *C. elegans* gene enrichment analysis tool, gave similar results [48]. Interestingly, WormCat analysis identified metabolism (sub-category ‘lipid metabolism’) as one of the major processes that was differentially influenced by *tcer-1* mutation and PA14 exposure (Fig. 1G). Amongst the lipid-metabolic factors impacted by these interventions were 5 of the 8 genes of the *‘lipl’* family. *lipl-1* and *lipl-2* were upregulated upon PA14 infection, and this upregulation was further enhanced in PA14-infected *tcer-1* mutants suggesting that these genes may contribute to the increased PA14 resistance observed in *tcer-1* mutants. We confirmed these observations using quantitative PCR (qPCR) analysis (Fig. 1H, I). Conversely, *lipl-3, lipl-4* and *lipl-5* were downregulated on PA14 and upregulated in *tcer-1* mutants on OP50 or PA14 (data not shown).

### TCER-1 differentially regulates *lipl-1* and *lipl-2* expression in adult tissues upon pathogen exposure

We next sought to identify the tissues and cells where *lipl-1* and *lipl-2* expression was being modulated. In a previous study, a 441bp *lipl-1* promoter-driven transgene was reported to be expressed in intestinal cells [37]. We created transgenic animals expressing mCherry driven by this promoter [*Plipl-1p(441bp):mCherry]*. Under normal conditions, *lipl-1* expression was observed at low levels in WT adults with fluorescence localized to the intestine and occasionally in the head (Fig. 2A-D). Within the transgenic population, fluorescence intensity ranged from quite dim, and primarily visible in the posterior intestine, to very bright throughout the intestine and neurons. A majority of animals exhibited an intermediate level of fluorescence (Figs. 2A-F and S2). The fraction of worms showing high intestinal fluorescence was increased significantly upon *tcer-1* RNAi as well as PA14 exposure (Fig. 2E), whereas, neuronal levels were not altered by either intervention (Fig. 2F). Pathogen exposure following *tcer-1* RNAi tripled and doubled the fraction of animals exhibiting high expression in the intestines and neurons, respectively (Fig. 2E, F) as predicted by the RNAseq data. We created an additional reporter strain driving mCherry under control of 1kb of *lipl-1* upstream region and found similar expression patterns (data not shown). To examine how these transcriptional changes translated into protein levels and localization *in vivo*, we generated a transgenic strain expressing a fluorescent-tagged LIPL-1::RFP driven by the 441bp promoter. LIPL-1::RFP expression showed similar spatial distribution as the mRNA in intestinal cells, but significantly lower levels. Expression in the head was punctate but inconsistent (Fig. 2G, K). Additionally, it was strongly localized to the coelomocytes (where mRNA signal was not observed), suggesting that LIPL-1 protein may be secreted into the pseudocoelomic cavity (Fig. 2G-J). *tcer-1* RNAi and PA14 exposure both appeared to induce modest and variable increases in RFP levels, paralleling the mRNA expression, though the highest changes were observed in coelomocytes (Fig. 2G-J).

**Figure 2:**
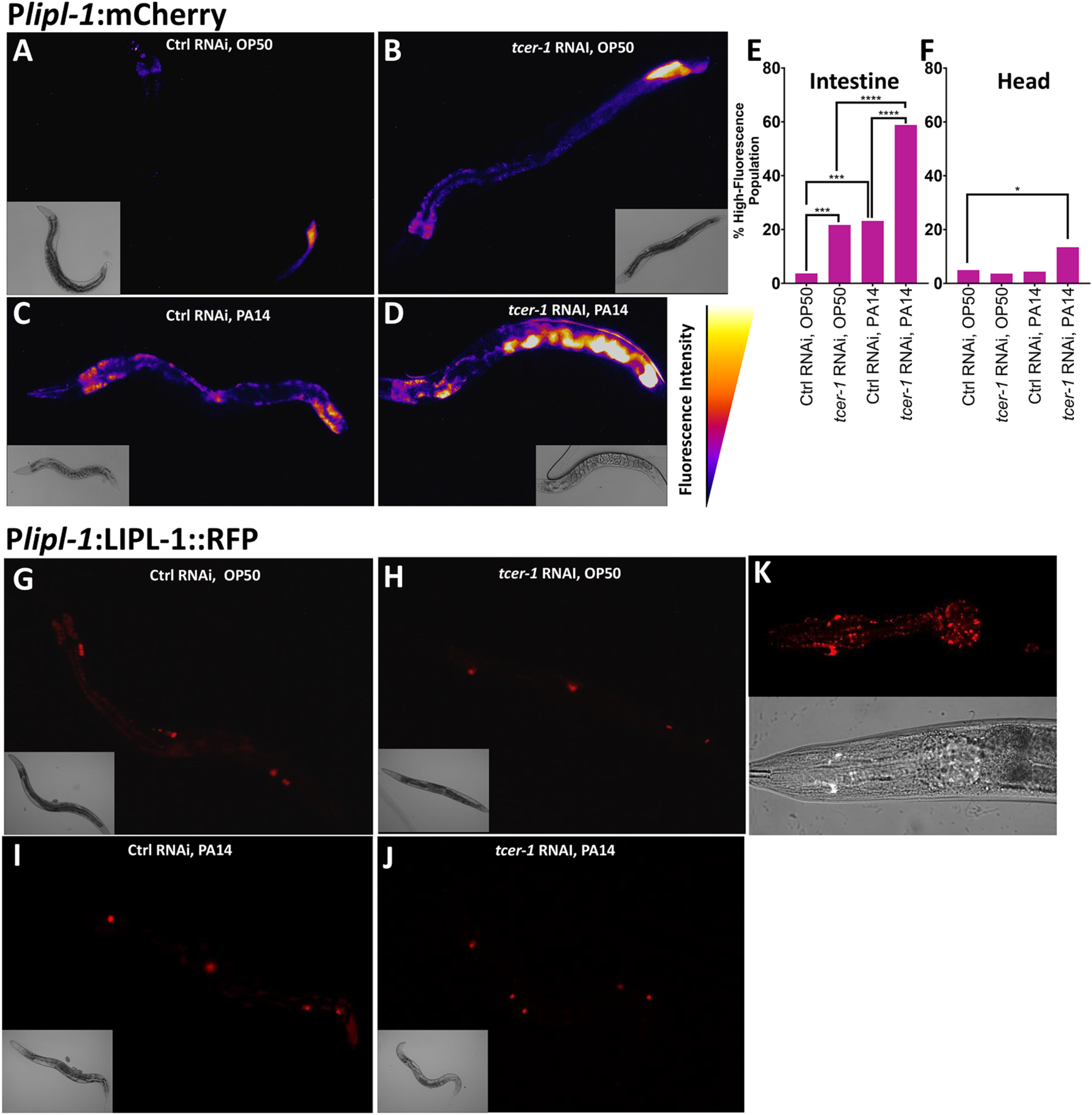
t*c*er*-1* inactivation and pathogen exposure induce *lipl-1* transcriptional upregulation. **A-F: *lipl-1* transcriptional changes visualized *in vivo* using *Plipl-1::*mCherry.** Animals raised on bacteria expressing control empty vector **(Ctrl, A, B)** or *tcer-1* dsRNA **(RNAi, C, D)** until pre-adult, L4 larval stage and transferred to plates seeded with *P. aeruginosa* PA14 **(PA14, C, D)** or *E. coli* OP50 **(OP50, A, B)** and incubated for 8 hours at 25^0^C. Expression visible in intestine and head, and in a normal adult population, varied from low levels to high or very high fluorescence intensities in intestine and head, respectively (categorization and quantification detailed in Fig. S2). **A-D** show representative images pseudocolored with ImageJ LUT Fire. Fraction of population with high or very high expression quantified in intestine **(E)** or head **(F)**. **E:** Control (Ctrl) RNAi OP50 (n=81, 3.7%), *tcer-1* RNAi, OP50 (n=83, 21.69%), Ctrl RNAi, PA14 (n=69, 23.19%), *tcer-1* RNAi, PA14 (n=68, 58.82%). **F:** Ctrl RNAi, OP50 (n=81, 10.52%), *tcer-1* RNAi, OP50 (n=83, 7.895%), Control RNAi PA14 (n=69, 7.895%), *tcer-1* RNAi, PA14 (n=67, 23.68%). Comparisons performed using two-tailed Fisher’s exact test on contingency tables of data from 3 pooled independent biological replicates. *p≤ 0.05, **p≤0.01, ***p≤0.001 and ****p≤0.0001. **G-K: LIPL-1 protein levels observed using translational reporter P*lipl-1*::LIPL-1::RFP.** Growth and exposure conditions similar to those in A-D. Representative images of Day 1 adults showing expression that was predominantly seen in coelomocytes (G-J) and rare animals with punctate expression in the head (K).

To examine *lipl-2* mRNA and protein localization *in vivo*, we similarly generated two endogenous promoter-driven mCherry-reporter strains (1kb and 1.5kb) and utilized a previously-published RFP-tagged strain (LIPL-2::RFP), respectively [39]. Both transcriptional reporters showed similar expression domains though the 1.5kb promoter-driven transgene showed more robust expression and was used to assess *lipl-2* regulation in this study. *lipl-2* transcription was also most prominent in anterior and posterior intestinal cells with few animals showing head fluorescence (Fig. 3A-D). *tcer-1* RNAi produced a modest elevation of intestinal reporter expression, whereas, PA14 exposure had no impact. Upon PA14 exposure in animals subjected to *tcer-1* RNAi, *lipl-2* expression in the intestine was strikingly elevated as compared to animals on control RNAi (Figs. 3E and S3A, B). No changes in expression were observed in the head under any condition. Interestingly, and unlike *lipl-1*, *lipl-2* exhibited age-related expression dynamics with higher expression observed during larval stages followed by a marked decrease in young adulthood and a subsequent induction with age, by Day 5, in posterior intestinal cells (Fig. S3C). LIPL-2::RFP signal was very low in normal adults, localized primarily to the intestine and coelomocytes (Fig. 3F-I) and with punctate, head fluorescence observed in a small number of animals; it was not visibly altered by PA14 or *tcer-1 lof* (Fig. 3J).

**Figure 3:**
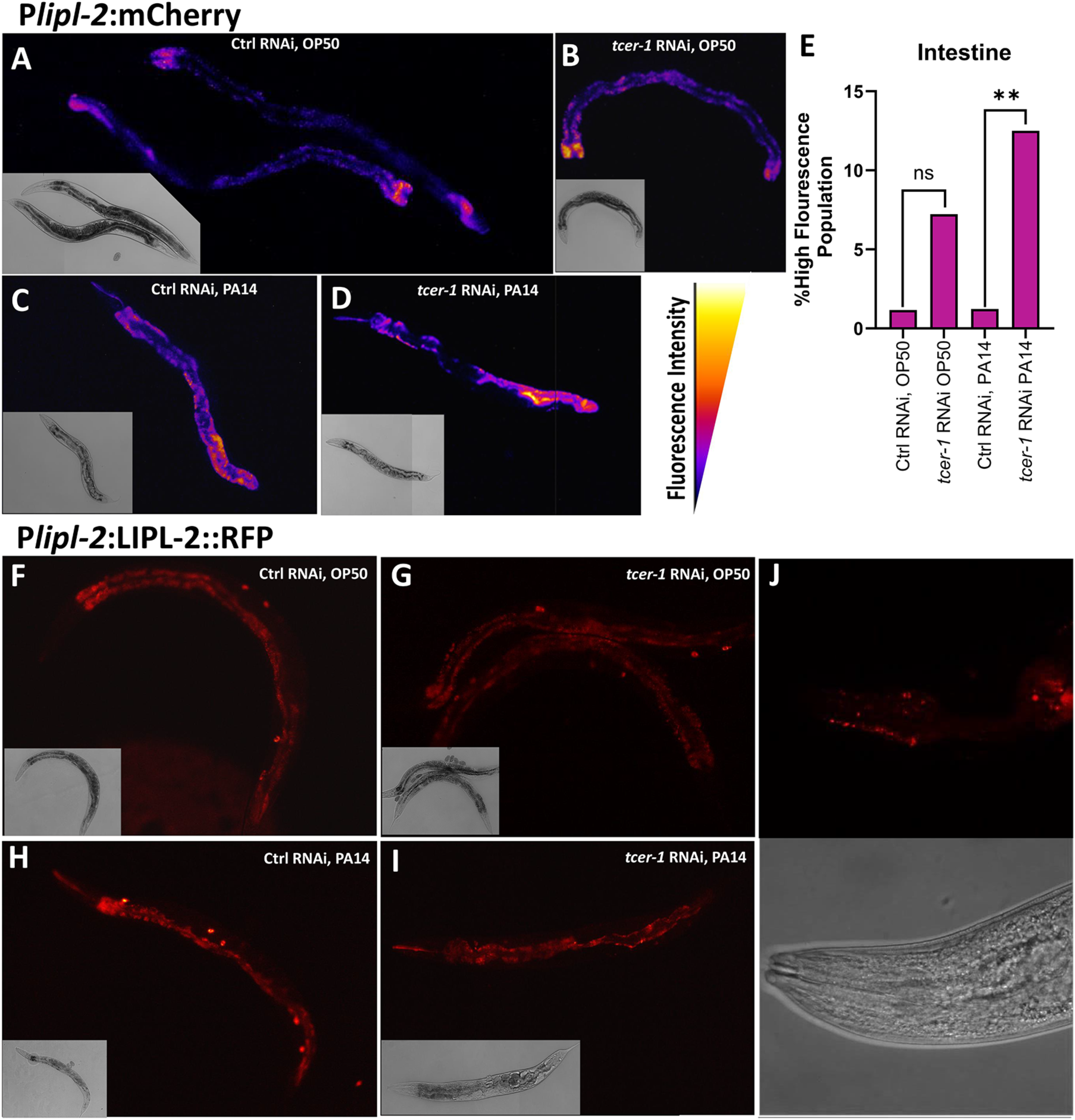
TCER-1 transcriptionally suppresses *lipl-2* and LIPL-2 protein levels are stringently controlled. *lipl-2* expression levels observed using the transcriptional reporter *Plipl-2::mCherry* **(A-E)** or translational reporter *Plipl-2:LIPL-2::RFP* (F-J). **A-E:** Animals expressing *Plipl-2:*mCherry were raised on bacteria expressing control empty vector (Ctrl) or *tcer-1* dsRNA (RNAi) until young adulthood, transferred to plates seeded with pathogen (PA14) or *E. coli* (OP50) bacteria and incubated for 8 hours at 25^0^C. Expression visible predominantly in the intestine and varied from low to high fluorescence intensities (detailed in Fig. S3). **A-D** show representative images pseudocolored with ImageJ LUT Fire. **E:** Fraction of populations showing high intestinal expression. Comparisons performed using two-tailed Fisher’s exact test on contingency tables of data from 3 pooled biological replicates of Control (Ctrl) RNAi OP50 (n=86, 1.16%), *tcer-1* RNAi, OP50 (n=83, 7.22%), Ctrl RNAi, PA14 (n=81, 1.23%), *tcer-1* RNAi, PA14 (n=72, 12.5%). *p≤ 0.05, **p≤0.01, ***p≤0.001 and ****p≤0.0001. **F-J: LIPL-1 protein levels observed using translational reporter P*lipl-1*::LIPL-1::RFP.** Growth and exposure conditions similar to those in A-D. Representative images of Day 1 adults pseudocolored with ImageJ LUT RedHOT showing very low expression predominantly in coelomocytes (with high intestinal autoflourescence in F-I). **J:** Rare animals with punctate expression in the head.

### LIPL-1, but not LIPL-2, is essential for the enhanced pathogen resilience of *tcer-1* mutants

We sought to determine the physiological impact of *lipl-1* and *lipl-2* on the immune response against PA14. Survival experiments using publicly-available, partial deletion mutants of the two genes yielded highly inconsistent results (data not shown). We used CRISPR-Cas9 to create complete deletion alleles of both genes (see Methods for details). We found that *lipl-1* was essential for the enhanced immune resistance of *tcer-1* mutants; in 4/5 trials, *lipl-1*;*tcer-1* mutants died significantly faster than *tcer-1* mutants and in 3 of these *tcer-1* mutants enhanced resistance was completely abolished (Fig. 4A, Table S3). *lipl-1* single mutants’ survival was not significantly different from that of WT animals (Fig. 4A, Table S3). Genes that enhance stress resistance, including immune stress, often increase longevity so we assessed the impact of *lipl-1* and *lipl-2* deletions on lifespan under normal conditions on an OP50 diet [49–54]. *lipl-1* deletion did not alter lifespan consistently, either alone or in a *tcer-1* mutant background (Fig. 4B, Table S4).

**Fig 4:**
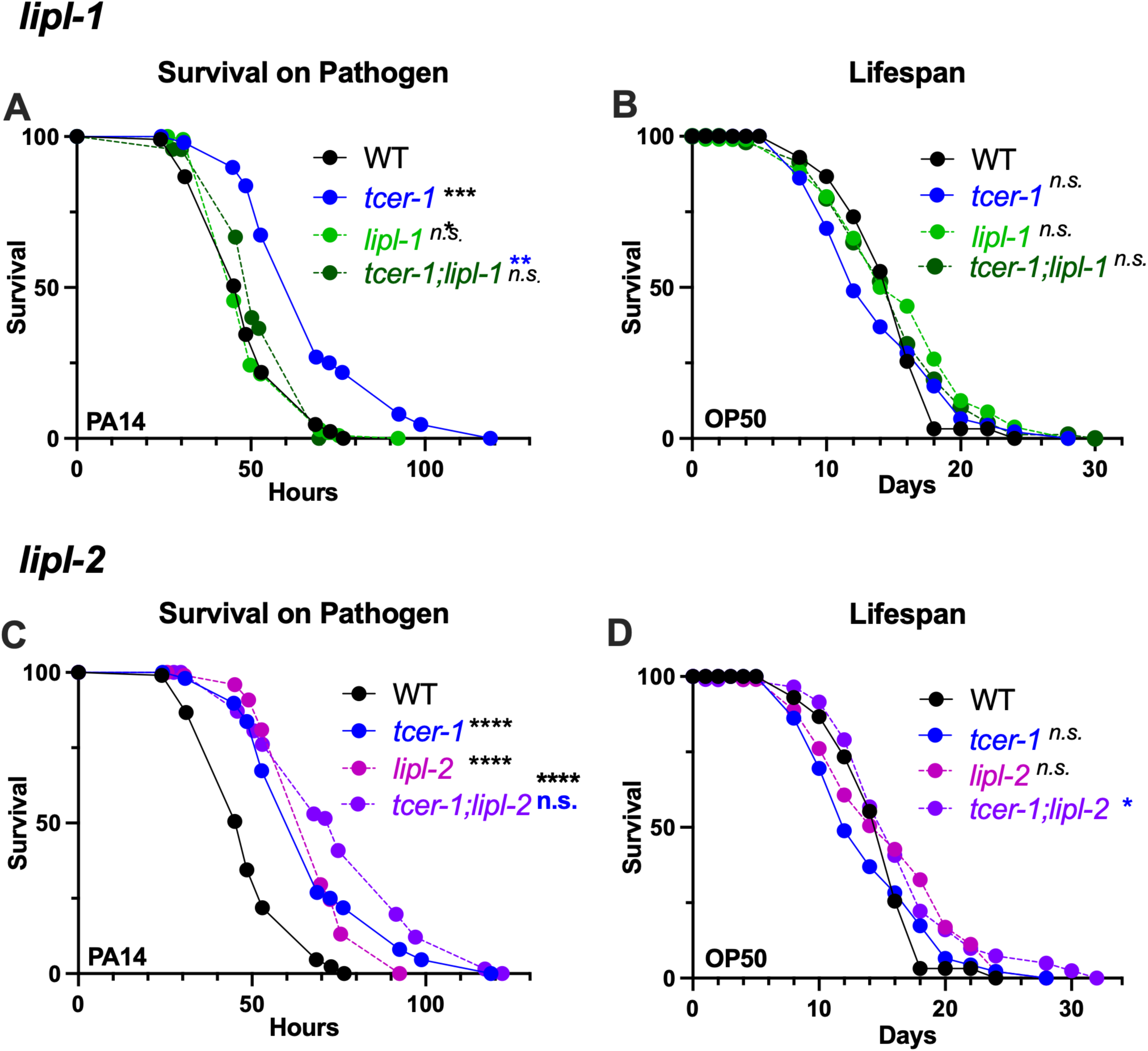
Enhanced immunity of *tcer-1* mutants is dependent upon *lipl-1* but not *lipl-2*. **A, B: Impact of *lipl-1* deletion on survival upon pathogen infection and lifespan.** Survival of wild type (WT, black), *tcer-1* (blue), *tcer-1;lipl-1* (dark green) and *lipl-1* (light green) raised on OP50 till L4 stage and exposed to PA14 **(A)** or retained on OP50 **(B). A:** WT (m= 51.13 ± 1.32, n= 86/113), *tcer-1* (m=70.61 ± 2.09, n=87/112), *lipl-1* (m=52.17 ± 1.1, n=102/134), *lipl-1;tcer-1* (dark green, m=56.04 ± 1.23, n=85/120). **B:** WT (m=14.96+-0.34, n=94/121), *tcer-1* (m=14.23 ± 0.48, n=92/120), *lipl-1* (m=15.79 ± 0.57, n=80/106), *lipl-1;tcer-1* (m=15.36 ± 0.51, n=77/120). **C, D: Impact of *lipl-2* deletion on survival upon pathogen infection and lifespan.** Survival of wild type (WT, black), *tcer-1* (blue), *tcer-1;lipl-2* (grape) and *lipl-2* (magenta) raised on OP50 till L4 stage and exposed to PA14 **(C)** or retained on OP50 **(D). C:** WT (m= 51.13 ± 1.32, n= 86/113), *tcer-1* (m=70.61 ± 2.09, n=87/112), *lipl-2* (m=70.64 ± 1.5, n=61/123), *tcer-1; lipl-2* (m=78.23 ± 2.66, n=66/124). **D)** Survival on OP50. WT (m=14.96 ± 0.34, n=94/121), *tcer-1* (m=14.23 ± 0.48, n=92/120), *lipl-2* (m= 15.93 ± 0.56, n= 89/113), *tcer-1; lipl-2* (m= 16.72 ± 0.58, n=81/103). Data from additional PA14 survival and lifespan trials in Table S3 and Table S4, respectively. Statistical significance was determined using log-rank Mantel Cox method and shown in each panel next to a given strain/condition. Asterisks are color coded to indicate the strain/condition being used for the comparison. p≤ 0.05(*), *p* < 0.01 (**), <0.001 (***), <0.0001 (****).

*lipl-2* inactivation had unexpected, contrasting effects on immune resistance and longevity compared to *lipl-1* mutants. The knockout did not suppress *tcer-1* mutants enhanced pathogen resistance and instead tended towards increasing it (Fig. 4C, TableS3). *tcer-1;lipl-2* mutants showed lifespan extension on a normal OP50 diet compared to *tcer-1* mutants (Fig. 4D, TableS4). *tcer-1* mutants exhibit increased survival upon exposure to other bacterial pathogens besides PA14, including Gram-positive *Staphylococcus aureus* [45]. We found that *lipl-1* and *lipl-2 lof* had similar impacts on the survival of *tcer-1* mutants and wild-type animals upon *S. aureus* exposure as well (Table S5). Together, these data suggest that *lipl-1* and *lipl-2* have distinct, context-dependent effects on immunity and longevity. *lipl-1* promotes immunity, especially upon *tcer-1* inactivation, but does not affect lifespan, whereas, *lipl-2* suppresses longevity, especially upon *tcer-1 lof*.

### LIPL-1 and LIPL-2 are essential for embryonic eggshell integrity and reproductive fitness

Since TCER-1 promotes fertility, we examined the effects of *lipl-1* and *lipl-2* inactivation on reproductive fitness in *C. elegans*. Expectedly, *tcer-1* mutants exhibited a marked reduction in the number of progeny produced (brood size) and deletion of *lipl-1* or *lipl-2* aggravated this phenotype significantly (Fig. 5A). Further, inactivation of either *lipl-1* or *lipl-2* induced sterility in *tcer-1* mutants (Fig. 5B). *tcer-1;lipl-1* and *tcer-1;lipl-2* mutants also displayed a reduction in the number of eggs that hatched into healthy larvae (hatching rate) (Fig. 5C). In *tcer-1*; *lipl-1 lipl-2* triple mutants, brood size was further diminished (Fig. 5A), whereas, the impacts on sterility and hatching-rate defects were not significant (Fig. 5B, C). In the presence of a functional TCER-1, deletion of *lipl-1* or *lipl-2* alone, or together, did not cause sterility or impair hatching rate (Fig. 5B, C). However, a modest reduction in brood size was manifested in both single mutants and this was not accentuated in the *lipl-1*;*lipl-2* double knockout (Fig. 5A).

**Fig 5:**
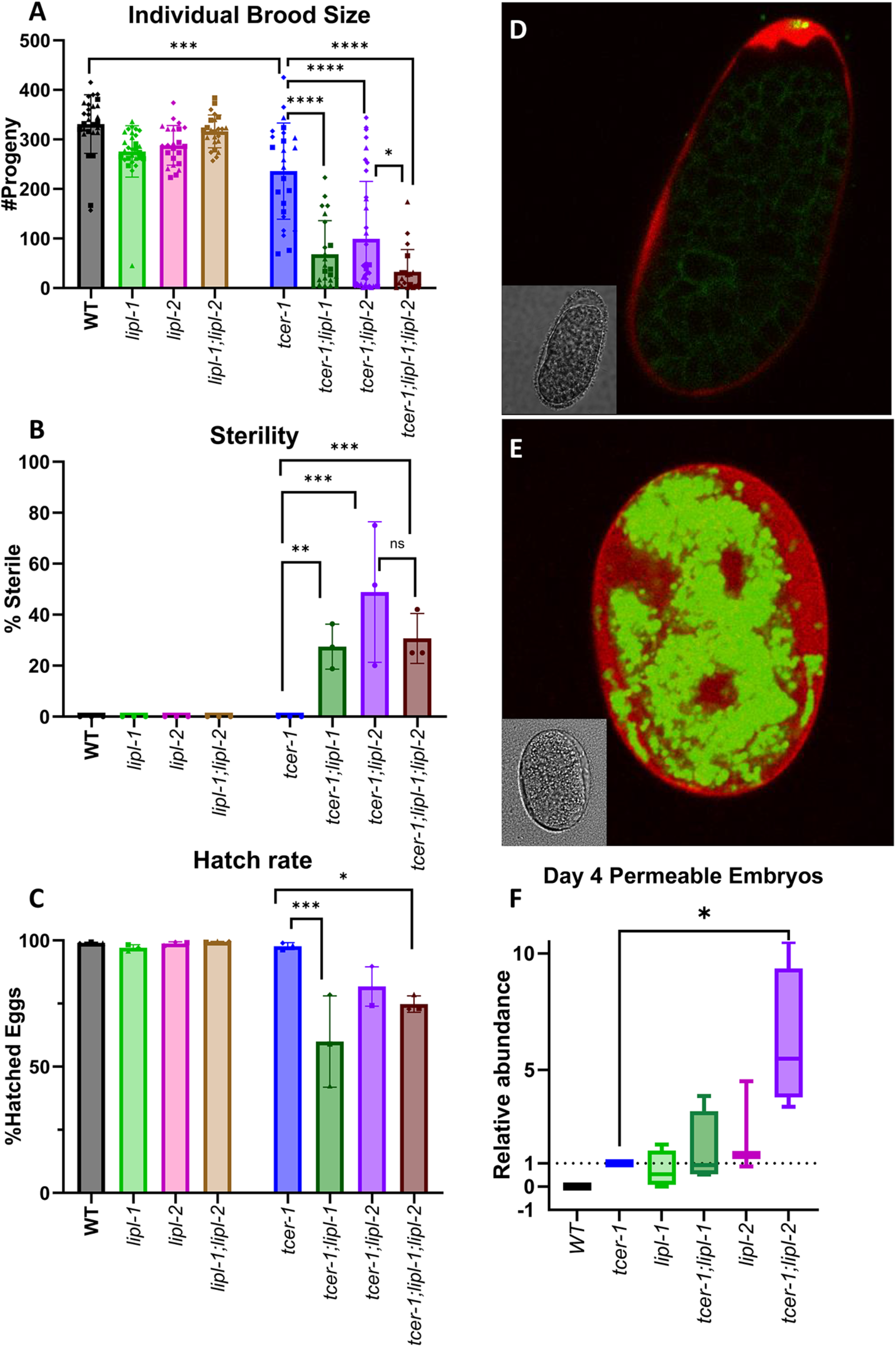
l*i*pl*-1* and *lipl-2 lof* result in decreased brood size, sterility, and embryonic inviability, particularly in combination with *tcer-1 lof.* **A-C: Comparisons of maternal reproductive success** in wild type (WT, black), *lipl-1* (light green), *lipl-2* (pink), *lipl-1 lipl-2* (light brown), *tcer-1* (blue), *tcer-1;lipl-1* (dark green), *tcer-1;lipl-2* (grape) and *tcer-1;lipl-1 lipl-2* (dark brown) strains. **A: Brood size:** Total number of live progeny counted per worm per strain. Each dot represents one worm from an aggregate of 3 independent trials. WT (n=28, m=330.9 ± 59.19), *lipl-1* (n=29, m=275.8 ± 51.97), *lipl-2* (n=23, m=288.2 ± 39.97), *lipl-1;lipl-2* (n=24, m=316.3 ± 33.23), *tcer-1* (n=25, m=236.2 ± 97.14) *tcer-1;lipl-1* (n=23, m=68.09 ± 68.12), *tcer-1;lipl-2* (n=36, m=99.31 ± 116.1), *tcer-1;lipl-1 lipl-2* (n=20, m=32.80 ± 44.87). **B: Sterility:** Percentage of animals which never produced a live progeny counted for each strain. WT (n=28, m=0), *lipl-1* (n=29, m=0), *lipl-2* (n=23, m=0), *lipl-1;lipl-2* (n=24, m=0), *tcer-1* (n=25, m=0), *tcer-1;lipl-1* (n=23, m=27.48 ± 8.82), *tcer-1;lipl-2* (n=36, m=48.87 ± 27.60), *tcer-1;lipl-1 lipl-2* (n=20, m=30.67 ± 9.81). **C: Hatch Rate:** calculated from [#progeny/(#progeny+ unhatched eggs)]. WT (m=99.01 ± 0.29), *lipl-1* (m=97.09 ± 1.24), *lipl-2* (m=98.81 ± 0.65), *lipl-1 lipl-2* (m=99.47 ± 0.15), *tcer-1* (m=97.75 ± 1.36), *tcer-1;lipl-1* (m=59.95 ± 18.07), *tcer-1;lipl-2* (m=81.75 ± 7.78), *tcer-1;lipl-1 lipl-2* (m=74.79 ± 3.24). Data obtained from 3 independent trials in all cases. **D-F: Embryonic eggshell defects induced upon maternal PA14 infection.** Representative images of embryos expressing *Pcpg-2::mCherry::CPG-2* and *pie-1p::GFP::PH PLC1delta1*, labeling eggshell chondroitin proteoglycan layer and embryonic plasma membrane, respectively, incubated with lipid-labeling dye, BODIPY. **D:** BODIPY is excluded in healthy embryo with intact lipid-permeability barrier (LBP) and CPG-2 is sequestered away from the embryo, in a characteristic ‘wavy’ pattern in the perivitelline space of the eggshell. **E:** Embryo with defective LPB exhibits widespread BODIPY staining, and mCherry::CPG-2 which freely diffuses between the outer eggshell and the embryo. **F:** Quantification of BODIPY-permeable embryos. Average fold-change of the fraction of BODIPY stained embryos laid by Day 4 mothers, normalized to *tcer-1* mutants. *tcer-1* (n=1964, m=0), WT (n=1930, m=0), *lipl-1* (n=2811, m=0.7153 ± 0.7889), *tcer-1;lipl-2* (n=1940, m=6.222 ± 3.030), *tcer-1;lipl-1* (n=2257, m=1.563 ± 1.587), *lipl-2* (n=2546, m=2.244 ± 1.980). Data obtained from 4 independent trials in all cases except *lipl-2* for which 3 trials were conducted. In A-C, statistical significance was calculated using one-way ANOVA with Tukey’s correction. In F, student’s t-test was used. p≤ 0.05(*), *p* <0.01 (**), <0.001 (***), <0.0001 (****).

As worms begin to age, they lay increasing numbers of unfertilized oocytes [33]. Previously, we had observed that *tcer-1* mutants produce larger number of unfertilized oocytes and from earlier ages than normal animals [33]. We asked if the *lipl-1* or *lipl-1* mutations were enhancing the hatching rate and brood size defects of *tcer-1* mutants by enhancing unfertilized oocyte production. However, no statistical difference between the oocyte production dynamics of the different mutant strains was observed (data not shown). Next, we examined the integrity of the embryonic eggshell, a protein and lipid rich structure critical for normal growth and development as it protects the embryo from mechanical and osmotic disruptions [55, 56]. In particular, the innermost layer of the eggshell, a fat-rich lipid-permeability barrier (LPB) serves as a deterrent against unregulated influx of external materials [55]. Hence, normal embryos are impermeable to lipid-staining dyes such as BODIPY or FM-64; when LPB is disrupted, the eggshell becomes permeable to these molecules and fluorescence can be observed within the embryonic body (Fig. 5D, E) [57, 58]. Embryonic porosity towards lipid dyes serves as a valuable measure of LPB and eggshell integrity. In WT animals, we did not find any embryos that were permeable to BODIPY when laid by young, Day 1 mothers, or even reproductively older Day 4 ones (Figs. 5F and S4). However, embryos laid by Day 4 mothers carrying *tcer-1* or either *‘lipl’* deletion produced a small but consistent fraction of eggs that accumulated BODIPY. The percent of porous eggs was not significantly increased in the *tcer-1;lipl-1* double mutants. In contrast, this population was enhanced by up to tenfold in *tcer-1;lipl-2* mutants (Fig. 5F). Indeed, *lipl-2* single mutants not only laid BODIPY-penetrable eggs on Day 4 of adulthood but also on Day 1 (Fig. S4).

Altogether, these data suggested that both lipases promote fertility in WT animals and *tcer-1* mutants, with *lipl-2* exerting a greater impact, and their absence leads to defective eggshell formation, reduced embryonic viability and fertility defects.

### TCER-1 and LIPL-2 Influence the Relative Distribution of Neutral Lipid Populations

To decipher the molecular changes brought about by TCER-1 on young animals’ lipid profile, and assess the contributions of LIPL-1 and LIPL-2, we performed high performance liquid chromatography coupled with tandem mass spectrometry (HPLC-MS/MS) on young Day 1 wild-type adults and age-matched *tcer-1*, *tcer-1;lipl-1* and *tcer-1;lipl-2* strains as well as *lipl-1* and *lipl-2* single mutants. HPLC-MS/MS allowed us to identify the impact of the genes on the abundance and relative distribution of major neutral lipid (NL) and phospholipid (PL) classes as well as individual lipid species within those classes. Within the global NL population, *tcer-1* mutants exhibited a significant shift in the relative abundance of the triacylglyceride (TAG) *vs.* diacylglyceride (DAG) levels. Compared to WT animals, their relative TAG content was increased by 15-20%, whereas, DAG levels were concomitantly decreased (Figs. 6 A, B and S5). Interestingly, *lipl-2* single mutants also showed a similar change in TAG:DAG distribution which was further aggravated in *tcer-1;lipl-2* double mutants. In contrast, *lipl-1* inactivation had little impact in either WT or *tcer-1* mutant backgrounds (Fig. 6A, B and Table S6) suggesting that, amongst the two lipases, *lipl-2*, but not *lipl-1* shaped the TAG:DAG distribution.

**Fig 6:**
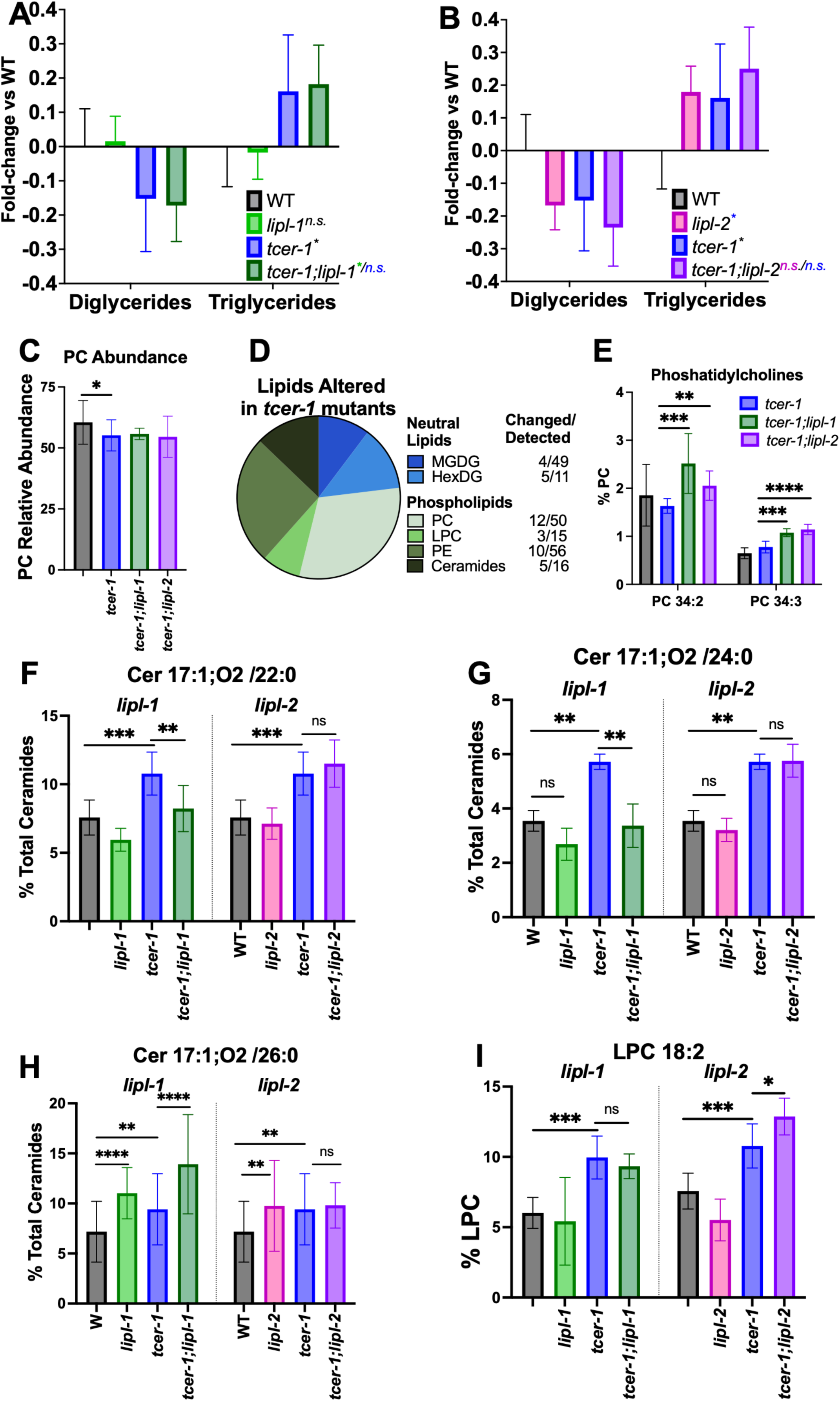
l*i*pl*-1* and *lipl-2* have both shared and distinct impacts on the lipidome of *tcer-1* mutants. **A-C: Effects of *lipl-1* (A) and *lipl-2* (B) loss on broad neutral lipid (NL) and phospholipid (PL) populations** in wild type (WT, black) and *tcer-1* mutant (blue) animals. Bars indicate the fold-change of each data point normalized to the WT average. Statistical significance calculated using mixed-effects analysis with Tukey’s correction. **C:** Relative abundance of overall phosphatidylcholine (PC) levels. **D:** Categorization of NL (top, blue) and PL (bottom, green) species whose abundance is altered in *tcer-1* mutants. For each class, number of species identified and those changed in *tcer-1* mutants shown in the table. **E:** Relative abundance of PC 34:2 and PC 34:3 compared to all PCs in different strains. **F-I: Distinct impacts of *lipl-1* (F-H) and *lipl-2* (I) on lipidome of *tcer-1* mutants. F-H**: Relative abundance of Cer17:1;O2/22:0 **(F)**, Cer 17:1;O2 /24:0 **(G)** and Cer 17:1;O2 /26:0 **(H)** compared to all ceramides. **I:** Relative abundance of lysophospatidyl cholines (LPC) 18:2 compared to all LPCs. Data derived from HPLC-MS/MS analysis of Day 1 young adult hermaphrodites isolated in 4-10 independent biological trials. Statistical significance was calculated using two-way ANOVA with Tukey’s correction. Asterisks indicate the statistical significance and their color the strain being used for the comparison. p≤ 0.05(*), *p*<0.01 (**), <0.001 (***), <0.0001 (****), n.s: not significant.

### LIPL-1 and LIPL-2 Impact the Levels of Phospholipid Species Implicated in Stress Resistance and Healthspan

In contrast to NLs, the relative abundance of the major PL classes was not significantly altered in *tcer-1* mutants except for a small but significant reduction in the levels of phosphatidylcholines (PC) (Fig. 6C). However, *tcer-1* mutants manifested changes in the abundance of specific PL species’ head groups. Of the 39 significantly altered lipid species in *tcer-1* mutants, 30 were PLs and 9 were NLs. Amongst PLs, PCs were the most affected class as 12 of 50 identified species were altered in *tcer-1* mutants-7 were decreased and 5 increased (Fig. 6D; Table S6). Besides PCs, phosphatidylethanolamines (PE) were the most disrupted (10 of 56 species identified) followed by lysophosphatidylcholines (LPCs) and ceramides (Cer) (Fig. 6D, Table S6). While neither *lipl-1* nor *lipl-2* deletion affected the total PC abundance in *tcer-1* mutants, both genes influenced the overall saturation of the associated fatty acid chains (Figs. S6, S7), and had shared impacts on the levels of two PCs-PC 34:2 and PC 34:3. Both species were significantly elevated in the *tcer-1;lipl-1* and *tcer-1;lipl-2* double mutants as compared to the *tcer-1* mutants (Fig. 6E, Table S6). *lipl-1* and *lipl-2* also exerted distinct effects on *tcer-1* mutants’ lipidome, most strikingly on the Cer and LPC compositions. *tcer-1* mutants showed increased levels of three Cer species, C17:1;O2/22:0, Cer 17:1;O2/24:0 and Cer 17:1;O2/26:0, and the elevation of all three species was significantly reduced in *tcer-1;lipl-1*, whereas, *tcer-1*;*lipl-2* mutants showed levels similar to *tcer-1* (Figs. 6F-H and S8, Tables S6 and S7). The effects on C17:1;O2/22:0 and Cer 17:1;O2 /24:0 paralleled the requirement for *lipl-1* for the enhanced PA14 resistance of *tcer-1* mutants, and pertinently C17:1O;2/22:0 has been implicated in stress resistance in *C. elegans* [59]. LIPL-2 had a similarly distinct impact on the abundance of LPC 18:2, which has been identified as a biomarker of healthspan and lifespan in human and *C. elegans*, respectively [60, 61]. LPC 18:2 levels were significantly enhanced in *tcer-1* mutants (Fig. 6I, TableS6). While *lipl-2* single mutants showed a small reduction compared to WT, in *tcer-1;lipl-2* mutants it was further elevated compared to *tcer-1* alone. *lipl-1* inactivation had no impact in either genetic background (Fig. 6I). The impact of *lipl-2* was specific to LPC 18: 2 as two other LPCs, 17:1 and 19:1, that were depleted in *tcer-1* mutants, were not changed by *lipl-2* deletion (Fig. S8, Table S6). *lipl-1* and *lipl-2* single mutants also manifested shared and distinct lipidomic changes compared to WT (Table S7). Altogether, these data showed that LIPL-1 and LIPL-2 had shared as well as discrete impacts on the lipidome, under normal conditions and upon *tcer-1* inactivation. Importantly, they revealed the influences of these genes on specific lipid moieties found previously to be associated with longevity and healthspan outcomes.

### Human Lysosomal Acid Lipase, LAL, Rescues the Immune Deficits Inflicted by Loss of *lipl-1*

*C. elegans* LIPLs are predicted to be orthologs of several human lipases, including LAL that is expressed in mammalian immune cells such as macrophages [62] and implicated in macrophage dysfunction, inflammatory signaling and lysosomal storage diseases [35, 63]. To assess a potential functional homology between *C. elegans* LIPL-1 and LAL, we used MosSCI technology to express hLAL in *C. elegans* and asked if the human protein could substitute for LIPL-1 in its immune function. We found that, upon PA14 infection, expression of hLAL in *tcer-1;lipl-1* mutants completely rescued their survival to *tcer-1* level (Fig. 7A, Table S8). In addition, transgenic strains expressing hLAL in somatic cells of WT *C. elegans* also survived substantially longer upon PA14 exposure (Fig. 7B, Table S8) suggesting a conserved immunity-promoting function.

**Fig 7:**
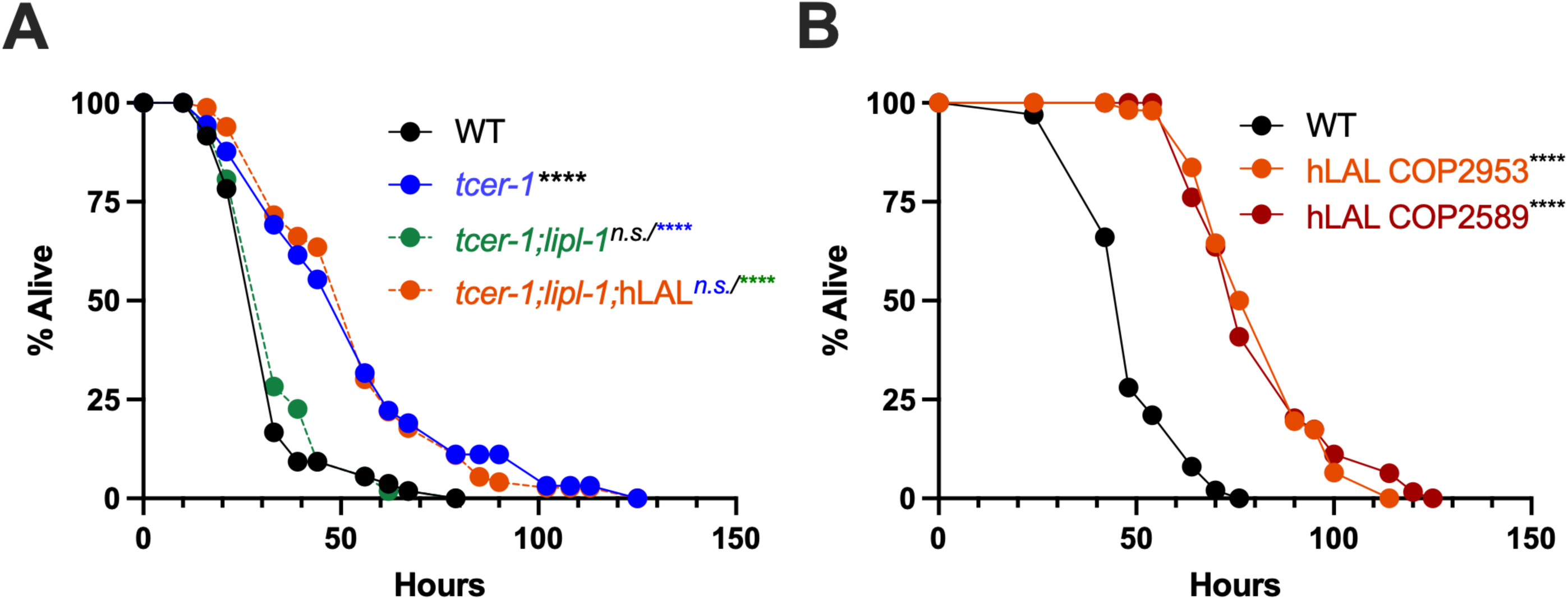
Human Lysosomal Acid Lipase (hLAL) rescues immune deficits induced by *lipl-1* mutation and enhances survival upon PA14 infection. **A: hLAL rescues survival of *tcer-1;lipl-1* mutants on PA14 exposure to *tcer-1* level.** Survival upon PA14 exposure from late-L4 stage onwards compared between wild type WT (black, m= 35.06 ± 1.91, n= 54/90), *tcer-1* (blue, m=54.8 ± 3.25, n= 63/90), *tcer-1;lipl-1* (green, m= 36.65 ± 1.79, n= 52/90), *tcer-1;lipl-1;*hLAL (green, m= 55.77 ± 2.53, n= 73/90). **B: hLAL expression in *C. elegans* enhances survival upon PA14 infection.** Survival upon PA14 exposure from late-L4 stage onwards compared between WT (m= 57.24 ± 2.01, n= 35/110) and two transgenic strains expressing hLAL broadly in somatic tissues. COP2983 (orange, m= 83.18 ± 2.22, n= 46/96) and COP2589 (red, m= 82.84 ± 2.22, n= 63/103). Statistical significance was calculated using the log-rank Mantel Cox method and is shown on each panel next to a given strain/condition with the color of the asterisk indicating strain being used for comparison. *p* < 0.01 (**), <0.001 (***), <0.0001 (****), n.s: not significant. Data from additional trials in Table S8.

## DISCUSSION

In this study, we identified *lipl-1* and *lipl-2* as mechanistic effectors of TCER-1, a transcriptional and splicing regulator that suppresses immunity and promotes fertility in *C. elegans*. Through transcriptomic, lipidomic, and genetic analyses, we show that these conserved lysosomal lipases perform distinct, context-specific roles in shaping immunity, reproduction, lipid profile, and lifespan. *lipl-1* enhances immune resistance in *tcer-1* mutants, whereas, *lipl-2* does not. Instead, *lipl-2* appears to limit longevity and immunity especially upon *tcer-1 lof*. Both lipases support fertility, especially embryonic integrity under infection stress, with *lipl-2* playing a more prominent role. Their lipidomic impacts are characterized by shared features well as influences on specific lipid species found to be correlated with stress resistance, lifespan and healthspan. Importantly, the human ortholog, LAL, rescues *lipl-1*-associated immune defects and improves survival upon infection suggesting potential evolutionary conservation of these molecular functions.

A central and unexpected finding of our study is the contrasting roles of *lipl-1* and *lipl-2* in immune defense. As per the RNA-seq prediction, *tcer-1* mutants showed elevated *lipl-1* during PA14 infection, relative to normal animals. Functionally, *lipl-1* was required for the enhanced pathogen resistance of *tcer-1* mutants but not wild-type animals, highlighting its context-specific role. Notably, *tcer-1* mutants accumulated the bioactive ceramide species, C17:1;O2/22:0, Cer 17:1;O2/24:0 and Cer 17:1;O2/26:0, and for the first two, the phenotype was reversed by *lipl-1* loss. Ceramides are essential membrane components that perform critical signaling functions in health and disease [64, 65]. In *C. elegans*, production of most ceramides is catalyzed by the ceramide synthase, *hyl-2,* and *hyl-2* mutants exhibit increased susceptibility to heat and oxidative stress [66, 67]. *hyl-2* was not identified as a DEG in our RNAseq so it is plausible that TCER-1 suppresses *lipl-1*-mediated ceramide production during PA14 infection; in its absence, *lipl-1* drives Cer 17:1;O2/22:0 accumulation to enhance pathogen resistance. This is consistent with prior evidence linking ceramide metabolism to stress tolerance and immunity [68]. Pertinently, ceramides are known to be generated by lysosomal lipases through the breakdown of complex glycosphingolipids and sphingomyelin [69].

The ability of the hLAL to substitute for *lipl-1* in promoting immune resistance is promising because of the known LAL roles in macrophage biology. In mammals, LAL regulates inflammation and is essential for M2 macrophage polarization, which supports anti-inflammatory responses and tissue repair [62]. Its deficiency causes lipid storage disorders such as Wolman’s disease and cholesterol ester storage disease (CESD) and leads to tissue-specific inflammation-parallels that reinforce a conserved role in immune regulation [35]. In mice, LAL deficiency leads to aberrant macrophage infiltration, while tissue-specific knockouts in the liver, intestine, and lung exhibit heightened inflammation in the affected tissues [70–72]. Given that M1/M2 macrophage polarization is critical for successful pregnancy [73], it will be interesting to test if hLAL also supports reproductive function.

*l*Our RNAseq data had led us to hypothesize that TCER-1 suppresses immunity by downregulating both *lipl-1* and *lipl-2* and had thus envisaged a pro-immunity function for both genes. Yet, *lipl-2* loss did not impair pathogen resistance and instead appeared to enhance post-infection survival and longevity in *tcer-1* mutants, underscoring that transcriptional changes do not always reflect functional impact. We also observed discrepancies between *in vivo* mRNA and protein reporters, with low mRNA and very low protein in adult tissues, suggesting tight post-transcriptional control and rapid protein turnover. The lipases’ protein reporters localized to coelomocytes despite intestinal transcription, implying secretion-consistent with prior findings on LIPL-1,3, and 5 [37, 38].Temperature also played a role: although *tcer-1 lof* consistently increased reporter expression in all conditions-matching gene expression data-the 25°C temperature required for PA14 virulence elevated reporter expression too, likely reflecting previously-described HSF-1–dependent heat shock response [74].

*lipl-2* supports reproductive health, particularly eggshell integrity during infection. Strikingly, the combined loss of *tcer-1* and *lipl-2* caused even young mothers to lay defective embryos. These phenotypes are noteworthy in light of the unique lipidomic changes we observed in *lipl-2;tcer-1* mutants. The abundance of 7 lipid species was altered in *lipl-2;tcer-1* mutants compared to *tcer-1* alone. Of these, elevation of three species-LPC 18:2, PC 32:1, and PC 32:2-has been linked to adverse pregnancy outcomes in women [75, 76]. Additionally, LPC 18:2 and PC 32:3, also elevated in *tcer-1;lipl-2*, are part of a panel of 8 serum biomarkers linked to human healthspan. In elderly individuals, higher LPC 18:2 levels predict slower gait decline, while PC 32:3 correlates with reduced gait speed. Although the effects of *lipl-2* are complex, these lipidomic parallels between *C. elegans* and human health metrics are compelling. Together with the rescue of *lipl-1 lof* immunity phenotypes by hLAL, they raise the enticing possibility that these genes may play conserved roles in the immune–reproductive axis.

## MATERIALS AND METHODS

### *C. elegans* and bacterial strains

All strains were maintained by standard techniques at 20°C or 15°C on nematode growth medium (NGM) plates seeded with an *E. coli* strain OP50. For experiments involving RNAi, NGM plates were supplemented with 1 mL 100 mg/mL Ampicillin and 1 mL 1M IPTG (Isopropyl β-D-1-thiogalactopyranoside) per liter of NGM. The main strains used in this study are listed in Supplementary Table S9.

### Generation of *lipl-1* and *lipl-2* Reporter Strains

*Translational Reporters:* The *Plipl-2*::LIPL-2::RFP construct was a gift from Dr. Abhinav Diwan. P*lipl-1*::LIPL-1::RFP was generated by amplifying 1,010 bp of the endogenous *lipl-1* promoter and entire CDS using primers which introduced Xma1 and Acc65i cut sites, and eliminated the CDS stop codon (Table 10). This was cloned into the P*lipl-2*::LIPL-2::RFP in PDG219 construct using restriction enzyme cloning to replace the *lipl-2* sequence. Transgenic strains were generated by injecting reporter constructs (25ng/ul) with a co-injection marker containing *myo-2*:GFP (15ng/ul) Strains were maintained by picking green fluorescent worms. *Transcriptional reporters:* For each construct, the selected region of the *lipl-1* (441bp or 994bp) or *lipl-2* (1000bp or 1500 bp) endogenous promoter was amplified using primers which added a homology region to the pAV1944 construct, containing *Pmyo-2*:mCherry. pAV1944 was linearized with Nhe1 and Sph1 restriction enzyme digestion to excise the *myo-2* promoter, and the final constructs were assembled using a gibson assembly kit (NEB E5520S) according to manufacturer directions. Transgenic strains were generated by injecting a *lipl-* reporter construct (50 ng/ul) along with a co-injection marker (*Pofm-1*:GFP or *Pmyo-2*:GFP, 15ng/ug). Strains were maintained by picking green fluorescent worms. Primers used in this study are listed in Supplementary Table 10.

### Generation of *lipl-1* and *lipl-2* deletion mutants

Strains were generated using the co-CRISPR method previously described[77]. Briefly, gRNAs and repair templates were designed to excise the entire CDS of *lipl-1* or *lipl-2* (Table 10). Injection mix (10ul volume: 24ug Cas9, 20ug tracrRNA (IDT 1072533), 4ug each target gRNA, 500ng target repair template, 2ug dpy-10 crRNA, 250ng dpy-10 repair template, IDT Nuclease-free Duplex Buffer to 10uL) was freshly prepared and spun for 20 minutes at 12,000g. Young D1 adults were injected in the gonad, and monitored for dumpy-roller progeny. F1 larvae from dpy/rol plates were screen via PCR to identify heterozygous CRISPR deletion alleles, which were outcrossed to N2 3x and homozygosed. Primers, gRNAs and repair templates used in this study are listed in Supplementary Table 10.

### Generation of hLAL strains

The transgenic strains COP2589 *{knuSi924 [pNU3447 (eft-3p::hLIPA::linker::wrmScarlet::3xFLAG::tbb-2u in cxTi10882, unc-119(+)) IV ; unc-119(ed3) III}* and COP2593 {*knuSi927 [pNU3447 (eft-3p::hLIPA::linker::wrmScarlet::3xFLAG::tbb-2u in cxTi10882, unc-119(+))] IV ; unc-119(ed3) III}* expressing hLAL using the pan-somatic promoter, *eft-3*, were created by InVivo Biosystems using the Mos1-mediated Singly Copy Insertions (MosSCI) method which enables integration of a transgene as a single copy at a designated *C. elegans* locus {Frøkjær-Jensen, 2015 #165}. The *unc-119* rescue cassette was used to bring the transgene into a target Mos1 locus on chromosome IV and create rescue of the *unc-119(ed3)* mutant allele. The Mos1 locus was selected for position-neutral effects and to avoid the gene-coding regions, introns and transcription factor binding sites. Transgene integration was confirmed by PCR.

### Flourescence Imaging and Quantification

*Transcriptional Reporters:* Transgenic animals were immobilized with 20 mM Levamisole, mounted on agar pads and imaged using a Leica DM5500B compound scope at 20x. Image acquisition was performed using LAS X software (Leica). Images were processed using ImageJ software. *Transcriptional Reporters:* Transgenic animals were immobilized with 20 mM Levamisole, mounted on agar pads and imaged using a Leica Stellaris 5 confocal with an integrated White Light Laser (WLL) at 20x and 63x. Image acquisition was performed using LAS X software (Leica). Images were processed using ImageJ software.

### Lifespan Assays

Lifespan experiments were performed as previously described [78]. All lifespan experiments were conducted at 20 °C on *E. coli* OP50 plates unless otherwise noted. Between 10-15 L4 hermaphrodites were transferred to each of ∼5−6 plates per experiment and observed at 24−48 h intervals to document live, dead or censored (animals that exploded, bagged or could not be located) animals. Animals were scored as dead when they failed to respond to gentle prodding with a platinum wire pick. Fertile strains were transferred every other day to fresh plates until progeny production ceased. The program Online Application of Survival Analysis 2 (OASIS 2) [79] was used for statistical analysis of both lifespan and pathogen stress assays. P-values were calculated using the log-rank (Mantel–Cox) test and subjected to multiplicity correction in experiments that involved more than two strains/conditions. Results were graphed using GraphPad Prism (Version 8).

### PA14-Infection Survival Assays

Pathogenic bacterial strain *Pseudomonas aeruginosa* (strain PA14) was streaked from frozen stocks onto Luria Bertani (LB) agar, incubated at 37 °C overnight and stored at 4 °C for a week or less. Single colonies from the streaked plates were inoculated and grown in King’s Broth to exponential growth phase, 6-12 hours at 37 °C with shaking. ∼20 µl of this broth culture was seeded onto slow killing (SK) plates (modified NGM plates containing 0.35% peptone instead of 0.25%) and incubated for 24 h at 37 °C. The plates were then left to sit at room temperature (RT) for 24 h prior to use. Between 30 and 40 L4 hermaphrodites per strain were transferred to each of ∼3-5 PA14 plates, incubated at 25 °C and monitored at 6−12 h intervals to account for live, dead or censored animals as described above. To rule out the impact of internal hatching on experimental outcomes, L4 larval stage animals were treated with 100 μg per ml of Fluoro Deoxy Uridine (fUDR) on NGM plates with OP50. Exposing *C. elegans* to this treatment for 24 h at 20^0^C before transferring to PA14 SK plates prevented the eggs from hatching. For experiments that also involved RNAi treatment, animals were grown to the L4 stage on standard RNAi plates seeded with *E. coli* HT115 strain carrying an empty vector control (pAD12 or L4440) or the relevant RNAi clone before transferring to PA14-seeded SK plates and assaying for survival at 25 °C. Kaplan−Meier analysis and statistics were performed as described above for lifespan assays.

### *S. aureus*-Infection Survival Assays

*S. aureus* killing assays were performed as previously described {Miller, 2015 #166}. *S. aureus* SH1000 {Horsburgh, 2002 #167} Δ*telA::Km^R^* (a gift from Dr. Lindsey Shaw) was grown overnight in tryptic soy broth (TSB) containing 50 μg/ml kanamycin (KAN). Overnight cultures were diluted 1:1 with TSB and 10 μl of the diluted culture was uniformly spread on the entire surface of 35 mm tryptic soy agar (TSA) plates containing 10 μg/ml KAN. Plates were incubated for 5 – 6 h at 37 °C, then stored overnight at 4 °C. Animals were treated with 100 μg/ml 5-fluoro-2′-deoxyuridine (FUDR) at L4 larval stage for ∼24 h at 15 °C - 20 °C before transfer to *S. aureus* plates. Three plates (technical replicates) were assayed for each strain in each biological replicate, with 20 - 40 animals per plate. Infection assays were carried out at 25 °C. Animals that crawled off the plate or died of bursting vulva were censored. Infection assays were carried out at least twice. Kaplan−Meier analysis and statistics were performed as described above for lifespan assays.

### Reproductive-Health Assays

Reproductive health was assessed using previously described methods. All experiments were conducted at 20°C and when matricide/bagging occurred the animal was censored from the experiment on that day. Individual synchronized L4 hermaphrodites were moved to fresh plates daily till the end of the reproductive phase (no progeny observed on a plate for a minimum of 2 days). Each day, once the parent was moved to a fresh plate, the older plate with eggs was stored at 20°C for ∼48 hours, and the number of hatched worms and eggs counted to calculate brood size. The above two parameters were used to determine viability (ratio of the total number of eggs laid by a hermaphrodite in its lifetime to total number of eggs that hatched). Similarly, the number of oocytes laid each day was counted to obtain oocyte number. These data were used to estimate the oocyte ratio (ratio of total number of oocytes laid by an animal to the total number of viable eggs it produced). Oocyte production span is the distribution of the oocyte ratio on a daily basis for the length of time that an animal lays any brood (eggs and oocytes combined). This parameter was used to assess premature oocyte production. The data presented here is obtained from aggregation of at least three independent trials, in each of which at least 10–15 animals per strain were examined.

### Embryo Permeability Assessment

For each genotype, approximately 20 gravid mothers were allowed to lay eggs overnight on 6cm plates. After 12hrs, the mothers were removed, and 200ul working solution of 20ug/mL BODIPY (Invitrogen D3922) in Egg Buffer [4 mM HEPES (pH 7.4), 94 mM NaCl, 32 mM KCl, 2.7 mM CaCl_2_ and 2.7 mM MgCl_2_] was added to flood each plate. After the liquid was completely absorbed, embryos were immediately counted and screened for fluorescence using scored using a Leica DM3000B microscope with a Lumen 200 metal halide fluorescence illuminator light box (Prior Scientific L200). For Day 4 experiments, the worms were transferred to fresh plates every day till Day 4 and then allowed to lay eggs before the same protocol was used.

### RNA-Sequencing and data analysis

RNA was isolated from 3 biological replicates of age-synchronized, Day 1 WT animals and CF2166 *tcer-1(tm1452)* mutants grown on OP50 till late L4 stage (∼48h), then transferred to PA14 plates (grown as described above) or allowed to continue on OP50. After 8hrs of exposure, worms were harvested for RNA isolation from approximately 3000 worms per strain.

Following 7 freeze thaw cycles, RNA was isolated using the Trizol method and for quality and quantity using the Agilent Tapestation and Qubit Fluorometry.

Sequencing libraries were prepared using the TruSeq stranded mRNA (PolyA+) kit and the samples were then subjected to 75 base pair paired-end sequencing on an Illumina NextSeq 500 sequencer at the Univ. of Pittsburgh Genomics Research Core. Sequencing data was analyzed using the CLC Genomics Workbench (Version 20.0.3) employing the RNA Seq pipeline. Differentially regulated genes were filtered for significant changes based on the criteria of >2 fold change in expression, P Value of <0.05 and a false discovery rate (FDR) of <0.05.

### Gene Ontology analyses

Genes that were differentially regulated in a statistically significant manner were classified into two groups as either up-regulated (UP) or down-regulated (DOWN) targets. These groups were analyzed for enrichment of gene classes based on Gene Ontology (GO) Terms using *C. elegans* centered publicly available online resources, Wormbase Gene Set Enrichment Analysis tool (https://wormbase.org/tools/enrichment/tea/tea.cgi) and WormCat (http://wormcat.com/) [80]. Representation Factor was calculated at http://nemates.org/MA/progs/overlap_stats.html.

### Lipidomics: Sample Collection

From each strain, ∼5,000 young Day 1 adults were collected at approximate onset of egg lay. To obtain these, approximately 600 gravid mothers were placed on 10cm plates seeded with OP50 and allowed to lay eggs for 5-8 hours, then moved to fresh plates to lay eggs overnight. The next day, all mothers were removed from overnight egg plates, and plates were subsequently monitored for onset of egg lay. WT, *lipl-1*, and *lipl-2* young adults were collected by washing plates with M9. *tcer-1*, *tcer-1;lipl-1*, and *tcer-1;lipl-2* populations exhibit delayed and unsynchronized development as well as high rates of infertility. To ensure that only fertile young adults were collected in these strains, once the population had abundant L4s, about 5,000 L4s were moved to a fresh 10cm plate. The next day, only individuals with eggs were included in sample collection. Due to the labor-intensive nature of this approach, collections were performed in two batches: (i) WT, *lipl-1*, *tcer-1*, *tcer-1;lipl-1* and [81] WT, *lipl-2*, *tcer-1*, *tcer-1;lipl-2.* Upon collection, worms were rinsed 2x with M9 and centrifugation (1000g x 2min). Final pellet was rinsed 1x with Millipore water and flash frozen. **Lipid Extraction:** Lipid extraction and detection of phospholipids (PL), neutral lipids (NL) and sphingolipids was carried out as described{Xatse, 2023 #168}. Briefly, for PLs, total lipids were extracted from nematodes strain using a chloroform/methanol (2:1 v/v) solvent system. A 1,2-diundecanoyl-sn-glyerco-3-phosphocholine standard was added for relative quantification prior to extraction (Avanti Polar Lipids). Total lipids were injected onto the LC-MS/MS system for the negative ion scanning mode analysis using an HPLC system (Dionex UHPLC UltiMate 3000) equipped with a C18 Hypersil Gold 2.1 × 50 mm, 1.9 μm column (25002-052130; Thermo Scientific) equipped with a 2.1 mm ID, 5 μm Drop-In guard cartridge (25005-012101; Thermo Scientific). Analysis was completed using a Q Exactive Orbitrap mass spectrometer coupled with a heated electrospray ionization source (Thermo Scientific). For extraction and detection of NLs, the same protocol was followed with minor changes. The standard for relative analysis was a triglyceride standard mix (GLC-406 Nu-Chek). Lipids were injected onto the HPLC-MS/MS system for the positive ion scanning mode analysis. For sphingolipid extraction and detection, a Ceramide/ Sphingoid Internal Standard Mixture I (Avanti Polar Lipids) was added to the nematode sample and total lipids extracted as previously mentioned using a chloroform:methanol mixture. Sphingolipids were separated and analyzed using the same instrument as PLs and NLs with the following modifications. 10 μl of the purified sphingolipids were injected onto the HPLC-MS/MS system for the positive ion scanning mode analysis. For MS analysis, the following parameters were used: the capillary temperature was set at 275◦C, the sheath gas flow rate was set at 45 units, the auxiliary gas flow was set at 10 units, the source voltage was 3.2 kV, and the AGC target was 106. Acquisition was carried out with full-scan data-dependent MS2 (ddMS2) mode. For MS1 profiling, scans were run at a resolution of 70k. MS2 analyses were performed using five scan events, where the top five ions were chosen from an initial MS1 scan. For fragmentation, a normalized collision energy of 50 was used. MS1 spectra were collected in profile mode, whereas MS2 spectra were collected in centroid mode. **Data Analysis:** Lipid analysis of the LC-MS/MS data was carried out using the software Lipid Data Analyzer (LDA) Version 2.8.1. LDA mass lists were generated for PLs and sphingolipids based on our previous studies{Dancy, 2015 #169}. LDA mass lists for NLs were created for all the neutral headgroups listed on LIPID MAPS. A relative quantification was used to compare between samples.

## Acknowledgements

The authors are grateful to members of the Ghazi lab and the Pittsburgh ‘Wormclub’ community for valuable inputs throughout this study. Thanks to Ho Yi-Mak (HKUST), John Murphy and Abhinav Dhawan (Washington University) for generously sharing strains. Some strains were provided by the CGC, which is funded by NIH Office of Research Infrastructure Programs (P40 OD010440). This work was supported by grants from the National Institutes of Health to AG (R01AG051659, 1R56AG066682, R01AI176326, R21AG083329) and JEI (R35GM149284, R21AI169842) and a Children’s Hospital of Pittsburgh Research Advisory Committee (RAC) graduate fellowship to LB.

## Author Contributions

AG conceptualized the study. AG, LB, CPO and JEI designed the experimental protocols. LB, FRGA, PS, DAB, BW, MC, GO, NN, MMC, JS and AG performed the experiments. AG, LB, FRGA, PS, JEI and CPO analyzed the data. LB and AG wrote the manuscript with input from the other authors.

---

**Supplementary Fig. S1:**
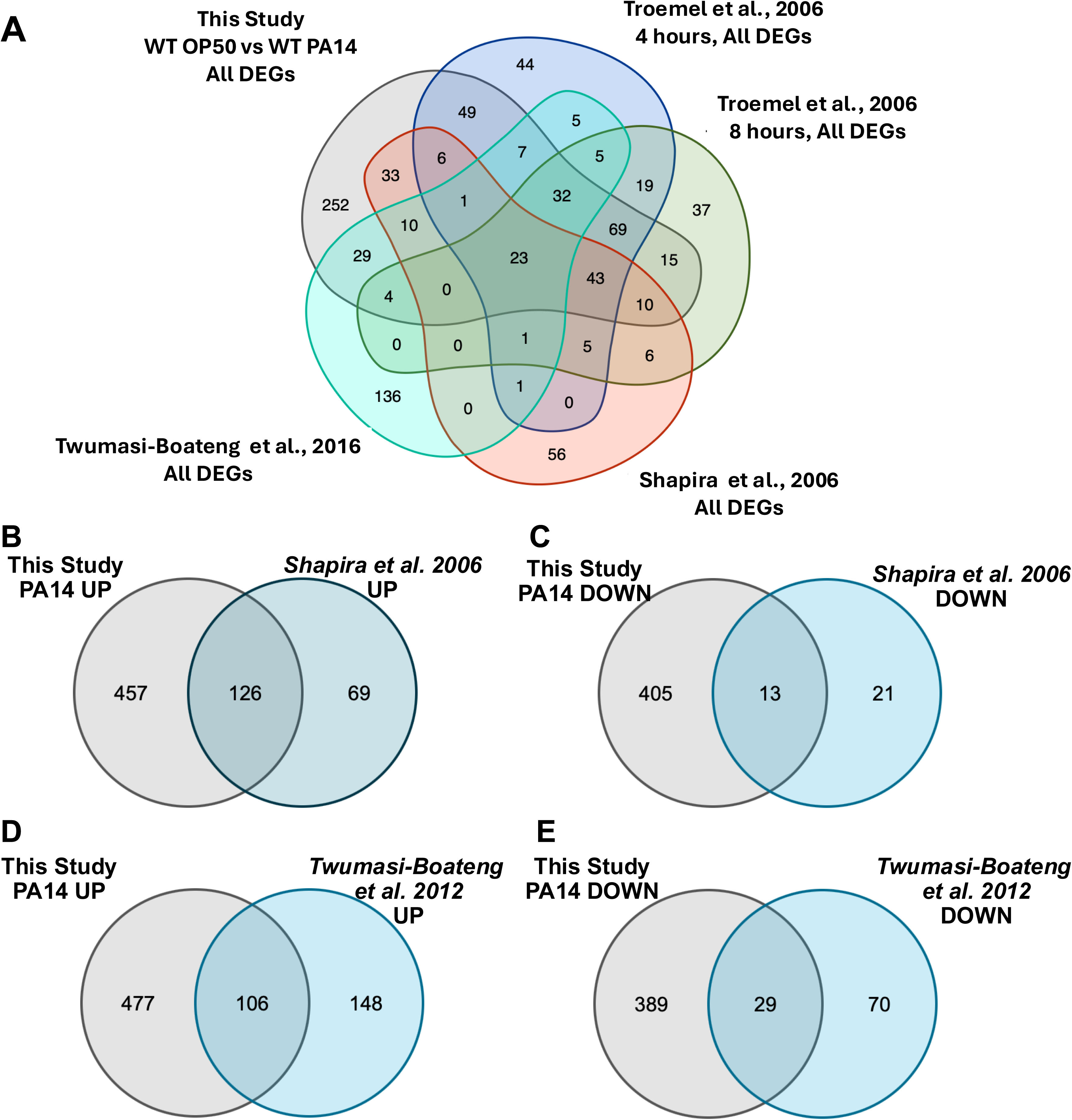
Overlap between PA14-induced differentially-expressed genes (DEGs) identified in this study and previously reported gene lists. **A:** All genes up- and down-regulated upon normal animals’ exposure to PA14 for 8 hours in this study compared with genes identified by Troemel et al. after 4 hours and 8 hours of exposure. **B-E:** Comparisons of DEGs upregulated **(B, D)** or downregulated **(C, E)** on PA14 with genes identified by Shapria et al., 2006 **(B, C)** and Twumasi-Boateng et al. 2012 **(D, E).** RF: Representation Factor. Statistical significance of overlap between gene sets calculated using hypergeometric probability formula with normal approximation (see Methods).

**Supplementary Fig. S2:**
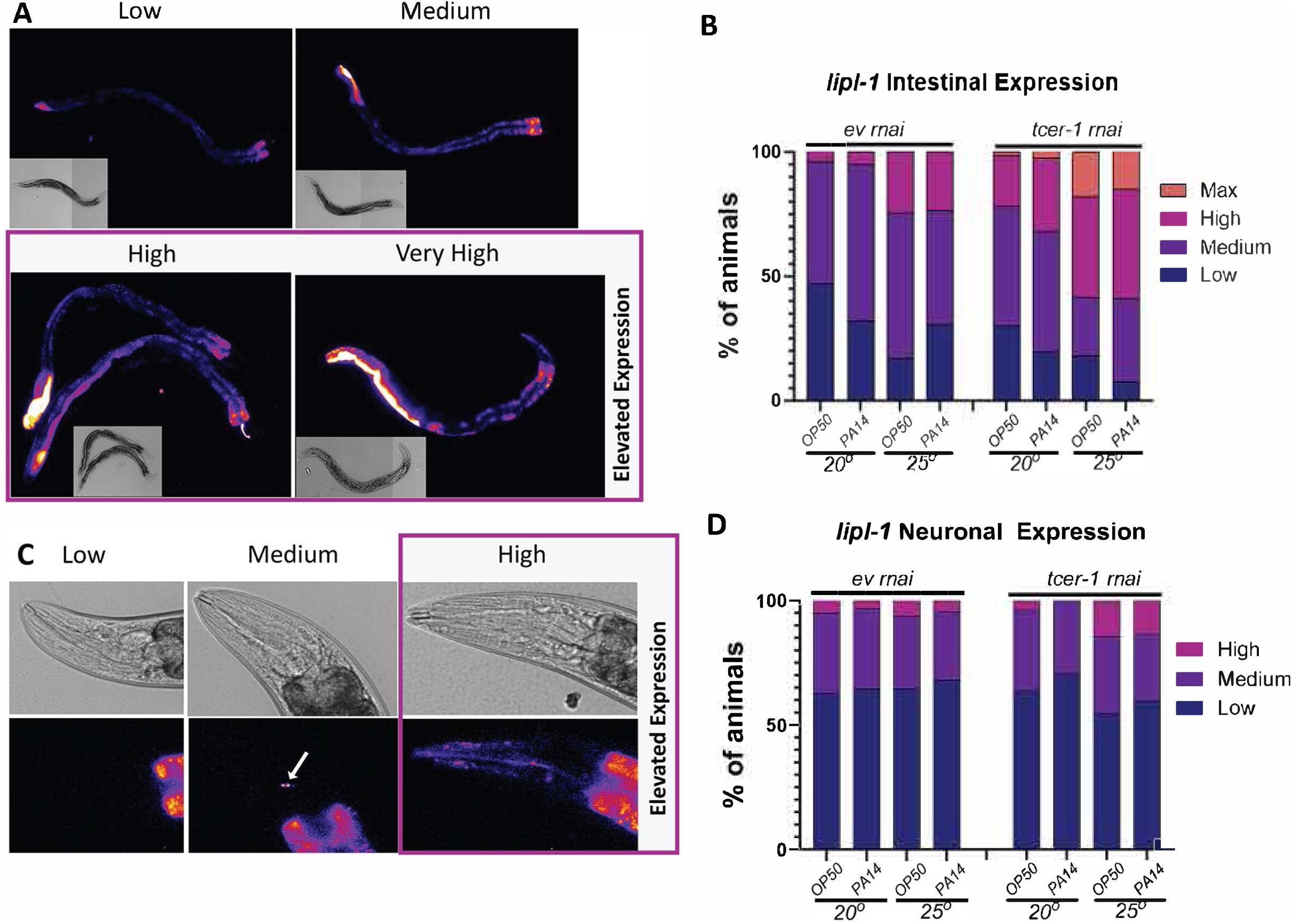
Categorical analysis of *Plipl-1::mCherry* expression. Flourescence levels quantified based on area and observable intensity variation (see Methods). **A, B: Intestinal Expression. A:** Representative images of each category pseudocolored with ImageJ LUT Fire. ***Low*-** dim fluorescence primarily visible in posterior and anterior intestine. No areas of high intensity, and posterior fluorescence limited to region near tail. ***Medium***-posterior fluorescence between vulva and tail. Some areas of intermediate intensity in anterior intensity. ***High***-posterior fluorescence uniformly extended up to the vulva or with multiple areas of bright intensity. ***Very High***– posterior intestine exhibited very bright fluorescence that extended into the anterior half beyond the vulva and intermediate fluorescence extended at least to the vulva. Purple boxes indicate categories quantified in Figure 2E, F. **B:** Quantification of percent of population in each category. Data from 3 pooled biological replicates. EV: Empty vector control. (ev RNAi till L4 then OP50, 20°C, n=81), (ev RNAi till L4 then PA14 20°C, n=62), (*tcer-1* RNAi till L4 then OP50, 20°C, n=83), *(tcer-1* RNAi till L4 then PA14 20°C, n=41), (ev RNAi till L4 then OP50, 25°C, n=82), (ev RNAi till L4 then PA14, 25°C, n=69), (*tcer-1* RNAi till L4 then OP50, 25°C, n=84), *(tcer-1* RNAi till L4 then PA14, 25°C, n=68) **C, D: Expression in Head Region. C:** Representative images of each category pseudocolored with ImageJ LUT Fire. ***Low*-** No flourescence visible. ***Medium*-** 1 or 2 puncta seen. ***High*-** More than 2 puncta, or diffuse, non-punctate expression. **D:** Quantification of data from 3 pooled biological replicates. (ev RNAi till L4 then OP50 20°C, n=81), (ev RNAi till L4 then PA14 20°C, n=62), (*tcer-1* RNAi till L4 then OP50 20°C, n=83), *(tcer-1* RNAi till L4 then PA14 20°C, n=41), (ev RNAi till L4 then OP50 25°C, n=82), (ev RNAi till L4 then PA14 25°C, n=69), (*tcer-1* RNAi OP50 25°C, n=84), *(tcer-1* RNAi PA14 25°C, n=67).

**Supplementary Fig. S3:**
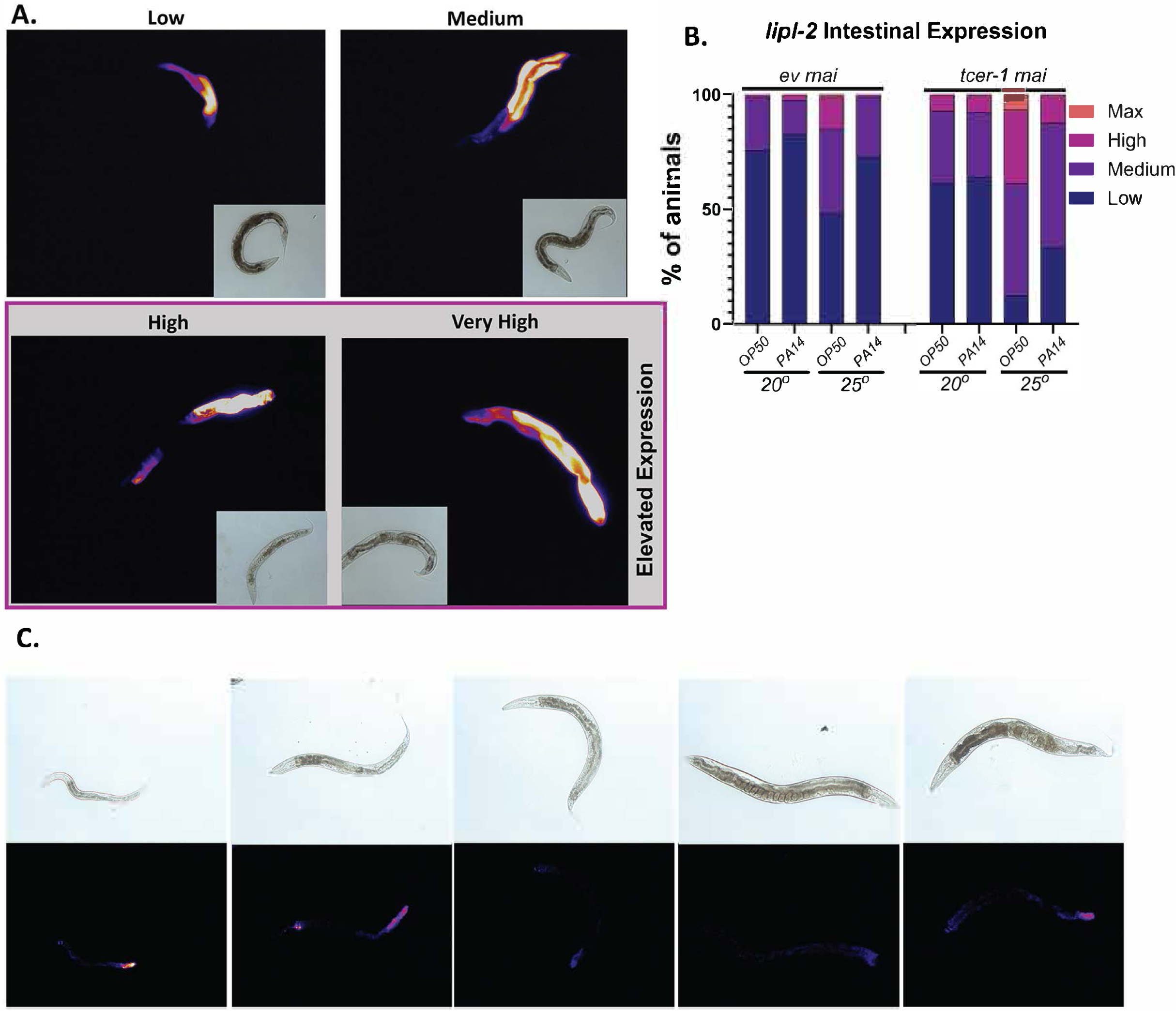
Categorical analysis of *Plipl-2::mCherry* expression. Expression levels in a population were variable and primarily observed in intestines. Flourescence quantified based on area and observable intensity variation (see Methods). **A, B: Intestinal Expression. A:** Representative images of each category pseudocolored with ImageJ LUT Fire. ***Low*-** dim fluorescence primarily visible in posterior intestine limited to region between vulva and tail. Brighter fluorescence, if present, was restricted to tail. ***Medium***-medium fluorescence in region between vulva and tail. Some area of increased intensity which extended past tail into anterior regions. ***High*** – broader posterior signal extending beyond vulva to anterior intestine with areas of bright intensity in posterior intestine. ***Very High***– Intense fluorescence extended from posterior intestine to anterior of vulva. Purple boxes indicate categories quantified in elevated expression analysis in Figure 2E, F. **B:** Quantification of percent of population in each category. Data from 3 pooled biological replicates. EV: Empty vectrol control. (ev RNAi till L4 then OP50 20°C, n=86), (ev RNAi till L4 then PA14 20°C, n=81), (*tcer-1* RNAi till L4 then OP50 20°C, n=83), *(tcer-1* RNAi PA14 till L4 then 25°C, n=78), (ev RNAi till L4 then OP50 25°C, n=87), (ev RNAi till L4 then PA14 25°C, n=81), (*tcer-1* RNAi till L4 then OP50 25°C, n=90), *(tcer-1* RNAi till L4 then PA14 25°C, n=72) **C: Expression dynamics across lifespan.** Representative images of *Plipl-2:*mCherry expression in (from left to right) young larvae, L4 larvae, young adult, Day 1 adult, and Day 5 adult worms. Pseudocolored in ImageJ with LUT Fire.

**Supplementary Figure S4:**
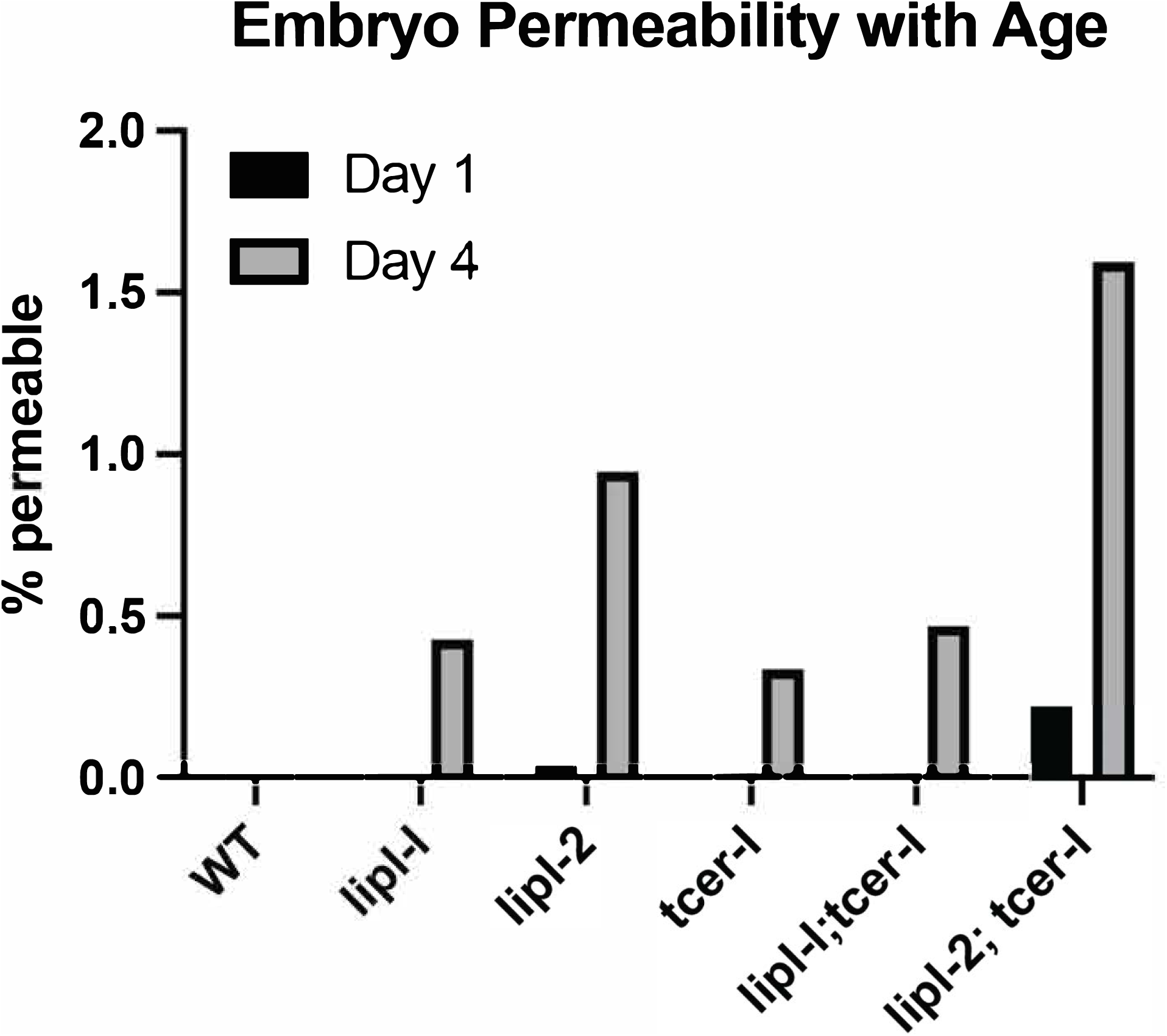
Loss of *tcer-1*, *lipl-1* and *lipl-2* causes embryonic eggshell defects. Longitudinal analysis of BODIPY-permeable eggs laid by Day 1 (D1) and Day 4 (D4) mothers by different strains. WT (D1 0, n=712; D4 0, n=276), *lipl-l* (D1 0, n=700; D4 0.4267, n=703), *lipl-2* (D1 0.0356, n=2806; D4 0.9451, n=529), *tcer-l* (D1 0, n=896; D4 0.3361, n=596), *tcer-1;lipl-1* (D1 0, n=1456; D4 0.4687, n=640), *tcer-1;lipl-2* (D1 0.2212, n=1356; D4 1.594, n=439).

**Supplementary Figure S5:**
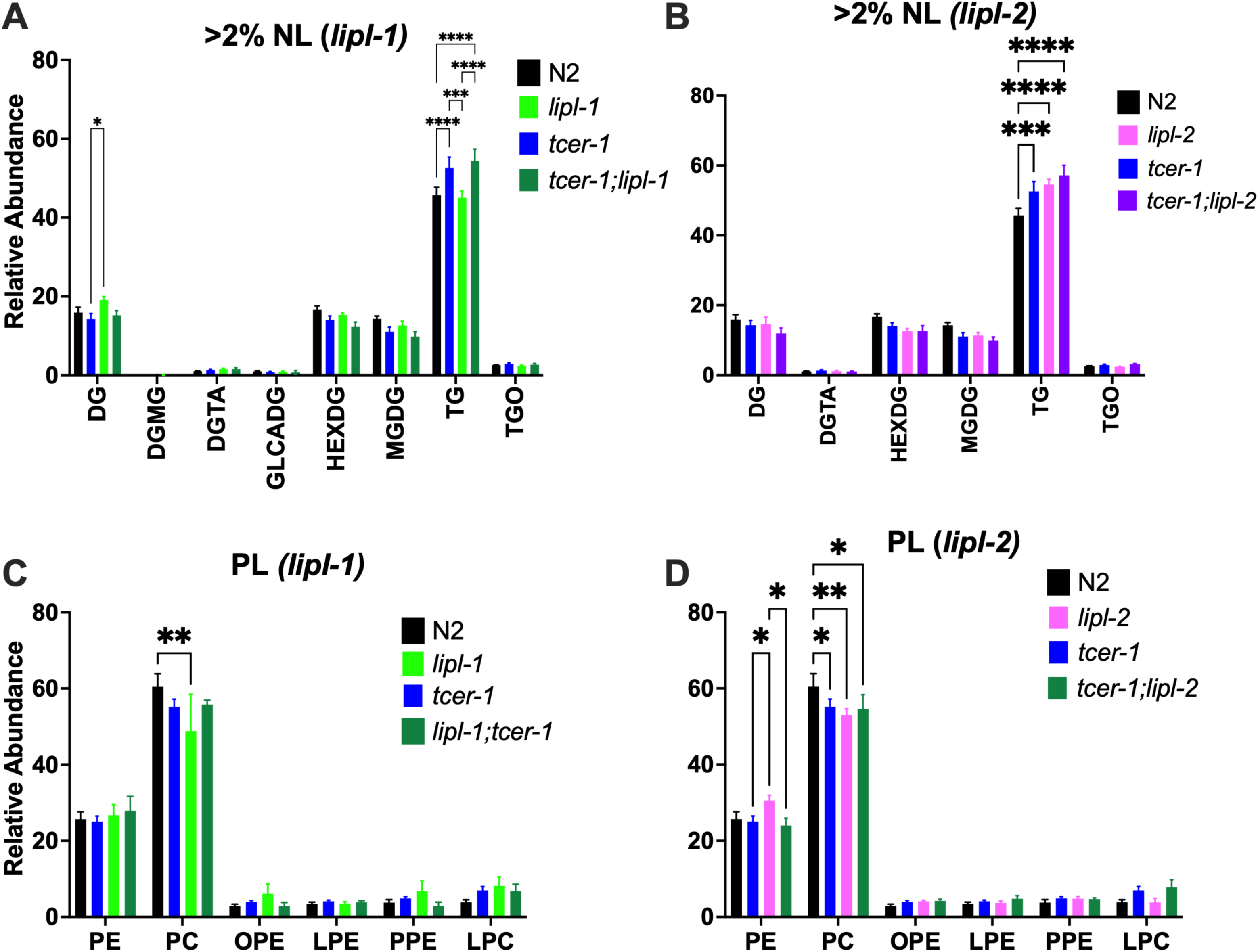
Impact of *lipl-1* and *lipl-2* deletions on relative abundance of neutral lipid (NL) and phospholipid (PL) classes. Lipids from gravid Day 1 adults were analyzed using HPLC-MS/MS. **A, B:** Impact of *lipl-1* (A) or *lipl-2* (B) inactivation on relative abundance of NLs with > 2% abundance in total NL population. **C, D:** Relative abundance of PL categories altered by *lipl-1* (C) or *lipl-2* (D) inactivation. Statistical significance was calculated using two-way ANOVA with Tukey’s correction, p≤ 0.05(*), *p* < 0.01 (**), <0.001 (***), <0.0001 (****).

**Supplementary Figure S6:**
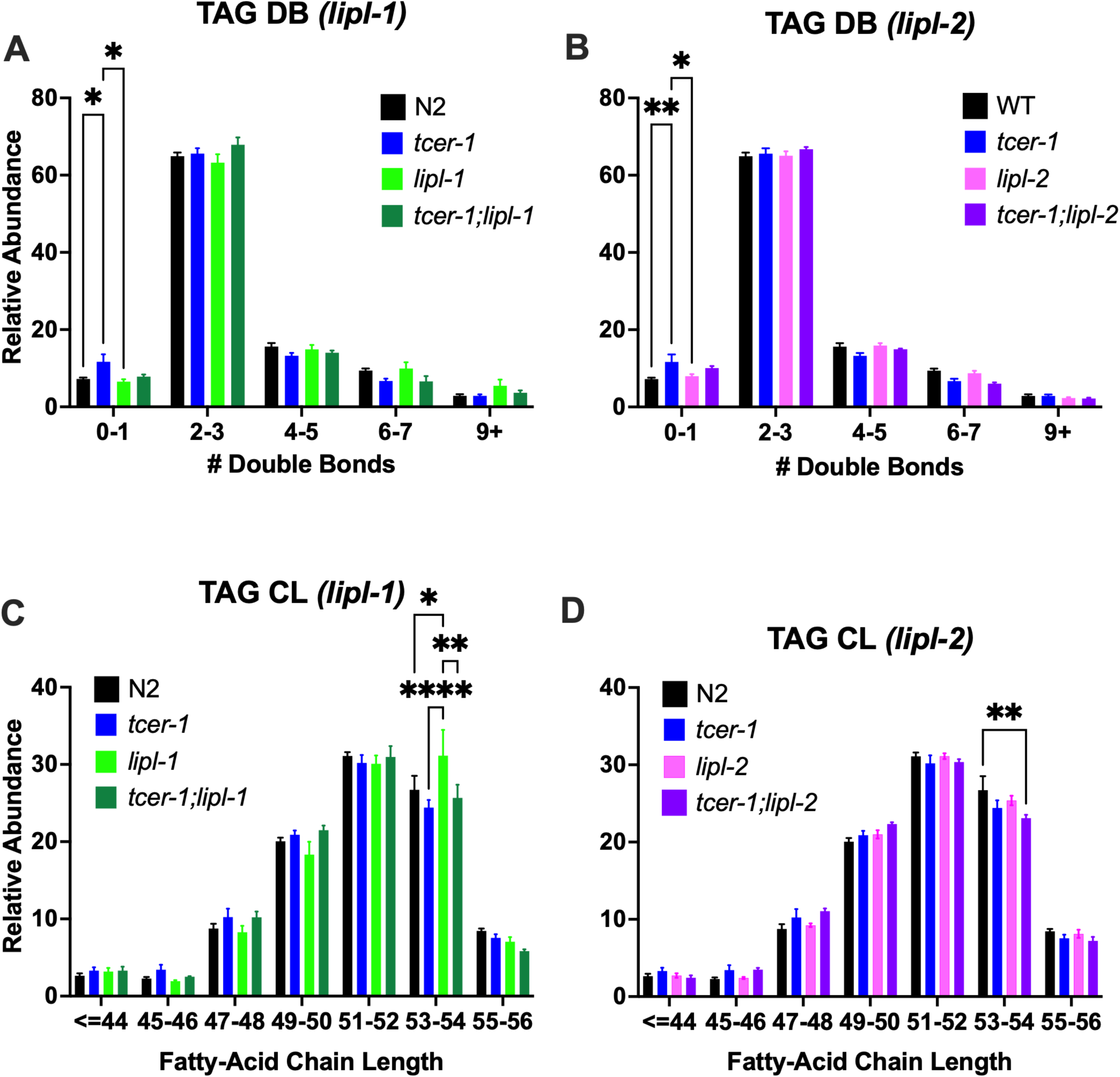
Impact of *lipl-1* and *lipl-2* deletions on saturation and fatty-acid chain length. **A, B:** Relative abundance of double bonds (DB) in triacylglycerides (TAG) altered by *lipl-1* (A) or *lipl-2* (B) inactivation. **C, D:** Relative abundance fatty-acid chain length (CL) of TAGs altered by *lipl-1* (C) or *lipl-2* (D) inactivation. Color key indicated on each panel. Statistical significance was calculated using two-way ANOVA with Tukey’s correction, p≤ 0.05(*), *p* < 0.01 (**), <0.001 (***), <0.0001 (****).

**Supplementary Figure S7:**
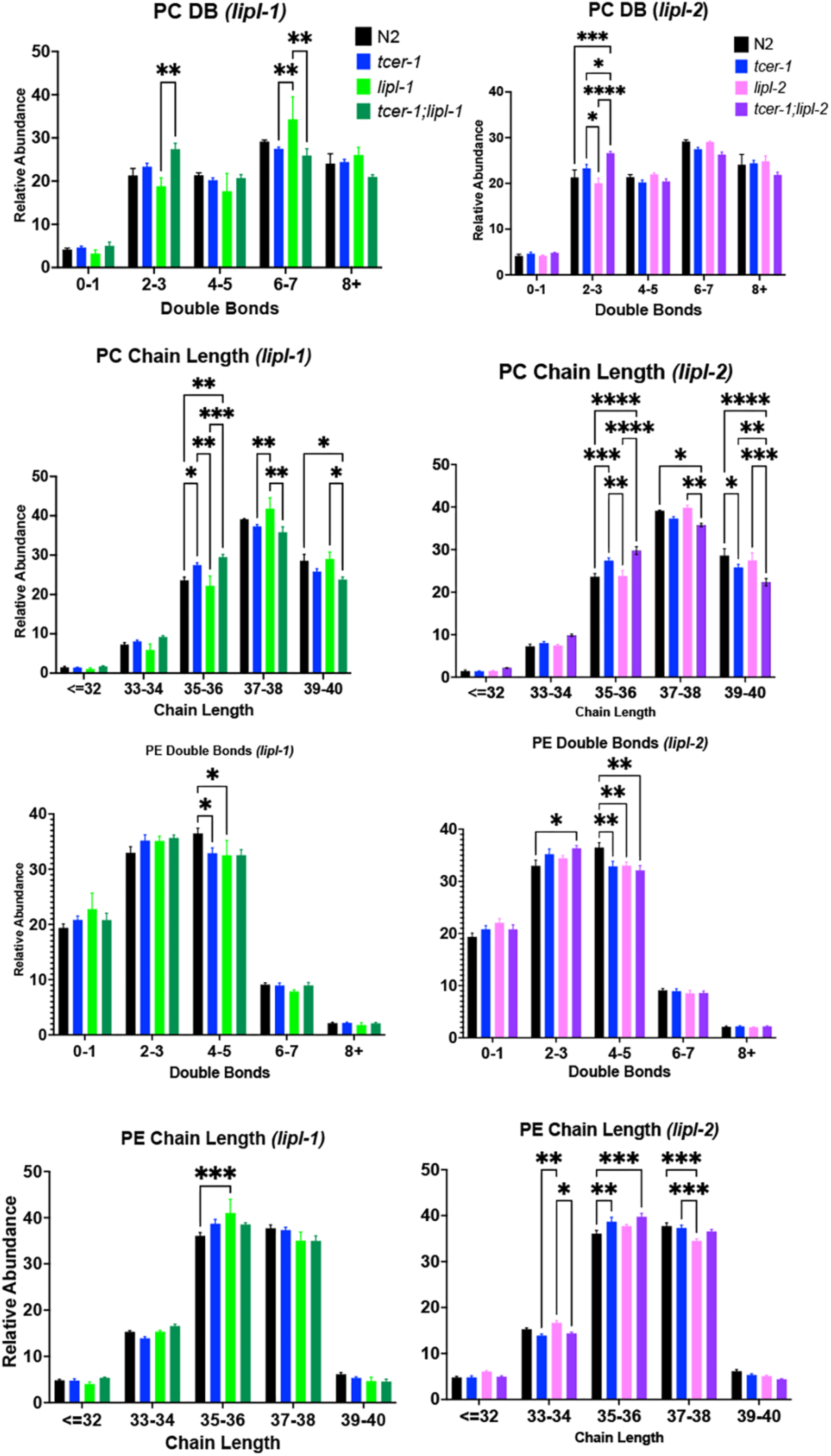
Impact of *lipl-1* and *lipl-2* deletions on saturation and fatty-acid chain length of phosphatidylcholine (PC) and phosphatidyl-ethanolamine (PE). **A, B:** Relative abundance of PC double bonds () altered by *lipl-1* (A) or *lipl-2* (B) inactivation. **C, D:** Relative abundance of PC chain length (CL) altered by *lipl-1* (C) or *lipl-2* (D) inactivation. **E, F:** Relative abundance of PE double bonds altered by *lipl-1* (E) or *lipl-2* (F) inactivation. **G, H:** Relative abundance of PE chain length altered by *lipl-1* (G) or *lipl-2* (H) inactivation. Color key of different strains shown at top. Statistical significance was calculated using two-way ANOVA with Tukey’s correction, p≤ 0.05(*), *p* < 0.01 (**), <0.001 (***), <0.0001 (****).

**Supplementary Figure S8:**
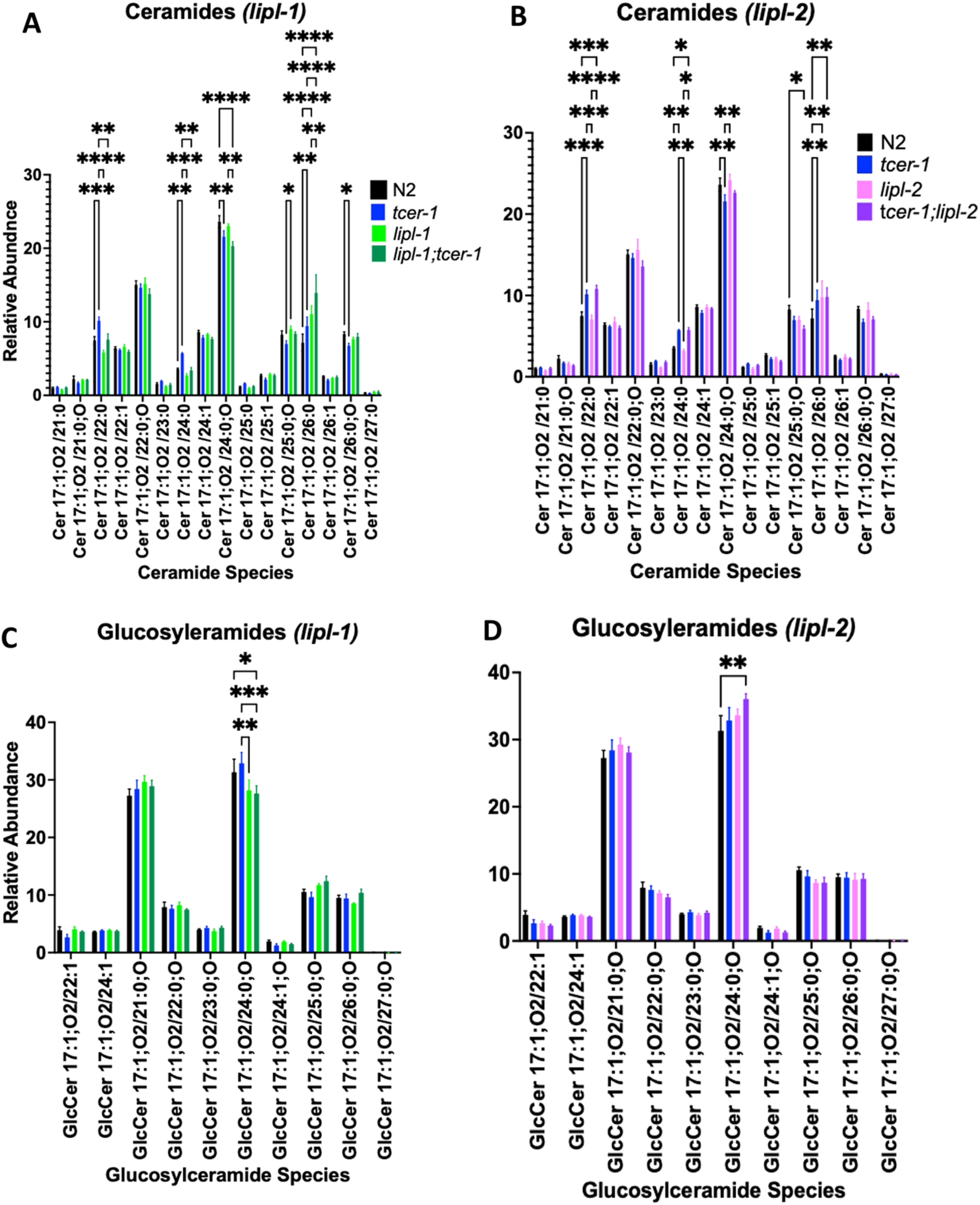
Impact of *lipl-1* and *lipl-2* deletions on glucosyl-ceramides (GlcCer) and ceramides (Cer). **A, B:** Relative abundance of GlcCers altered by *lipl-1* (A) or *lipl-2* (B) inactivation. **C, D:** Relative Abundance of Cers altered by *lipl-1* (C) or *lipl-2* (D) inactivation. Color key of different strains shown at top. Statistical significance was calculated using two-way ANOVA with Tukey’s correction, p≤ 0.05 (*), *p* < 0.01 (**), <0.001 (***), <0.0001 (****).

## List of Supplementary Tables

**Table S1:** Gene lists and GO-term analyses of differentially expressed genes identified by RNAseq in this study.

**Table S2:** Overlaps between PA14-induced genes identified in this study and previously reported PA14-driven genes.

**Table S3:** Impact of *lipl-1* and *lipl-2* null mutants on survival upon *P. aeruginosa* PA14 infection.

| ***lipl-1* Strains** |  |  |  |  |  |  |  |
| --- | --- | --- | --- | --- | --- | --- | --- |
| Strain | Background Genotype | Trial 1 |  |  | Bonferroni P-value |  |  |
|  |  | n = obs/ total | Mean | SE ^ | P (vs N2) | P (vs *tcer-1*) | P (vs *lipl-1*) |
| N2 | WT | 75/98 | 49.72 | 0.86 |  |  |  |
| CF2166 | *tcer-1* | 91/147 | 64.03 | 1.57 | **0.00000002** |  |  |
| AGP347 | *lipl-1* | 59/119 | 52.07 | 1.4 | **0.0184** |  |  |
| AGP354 | *tcer-1;lipl-1* | 75/135 | 56.32 | 1.57 | **0** | 0.1229 |  |
| Trial 2 |  |  |  |  | Bonferroni P-value |  |  |
| N2 | WT | 56/95 | 52.88 | 1.35 |  |  |  |
| CF2166 | *tcer-1* | 68/128 | 63.69 | 1.63 | **0** |  |  |
| AGP347 | *lipl-1* | 78/160 | 49.68 | 0.87 | 0.7954 | **0.000000014** |  |
| AGP354 | *tcer-1;lipl-1* | 101/126 | 48.17 | 0.8 | **1** | **0** |  |
| Trial 3 |  |  |  |  | Bonferroni P-value |  |  |
| N2 | WT | 41/50 | 56.64 | 2.05 |  |  |  |
| CF2166 | *tcer-1* | 100/120 | 71.59 | 2.04 | **0.0001** |  |  |
| AGP347 | *lipl-1* | 104/110 | 54.06 | 1.14 | 0.152 |  |  |
| AGP354 | *tcer-1;lipl-1* | 51/64 | 58.17 | 1.9 | **1** | **0.0002** |  |
| Trial 4 |  |  |  |  | Bonferroni P-value |  |  |
| N2 | WT | 76/93 | 52.58 | 1.41 |  |  |  |
| CF2166 | *tcer-1* | 68/100 | 66.56 | 1.55 | **0.00000015** |  |  |
| AGP347 | *lipl-1* | 77/94 | 56.29 | 1.19 | **1** |  |  |
| AGP354 | *tcer-1;lipl-1* | 73/95 | 59.76 | 1.34 | 0.1131 | **0.0001** |  |
| ***lipl-2* Strains** |  |  |  |  |  |  |  |
| Strain | Background Genotype | Trial 5 |  |  | Bonferroni P-value |  |  |
|  |  | n = obs/ total | Mean | SE ^ | P (vs N2) | P (vs *tcer-1*) | P (vs *lipl-2*) |
| N2 | WT | 102/119 | 59.27 | 1.21 |  |  |  |
| AGP364a | *lipl-2* | 117/126 | 60.74 | 1.47 | **1** |  |  |
| AGP336a | *tcer-1* | 98/123 | 67.44 | 1.51 | **0** |  |  |
| AGP358 | *tcer-1;lipl-2* | 84/127 | 75.27 | 1.85 |  | **0.0045** |  |
| AGP360a | *tcer-1; lipl-1 lipl-2* | 124/138 | 55.56 | 1.34 |  | **0** | **0** |
| Trial 6 |  |  |  |  | Bonferroni P-value |  |  |
| N2 | WT | 113/148 | 63.79 | 1.25 |  |  |  |
| AGP364a | *lipl-2* | 107/123 | 72.55 | 2.15 | 0.1846 |  |  |
| AGP336a | *tcer-1* | 123/134 | 91.69 | 2.25 | **0** |  |  |
| AGP358 | *tcer-1;lipl-2* | 100/129 | 95.53 | 3.07 |  | **1** |  |
| AGP360a | *tcer-1; lipl-1 lipl-2* | 107/140 | 67.58 | 1.71 |  | **0** | **0** |
| Trial 7 |  |  |  |  | Bonferroni P-value |  |  |
|  | WT | 86/113 | 51.13 | 1.32 |  |  | vs. *lipl-1* /vs. *lipl-2* |
| AGP347 | *lipl-1* | 102/134 | 52.17 | 1.1 | **1** |  |  |
| AGP364a | *lipl-2* | 61/123 | 70.64 | 1.5 | **0** |  |  |
| AGP357a | *lipl-1 lipl-2* | 83/117 | 51.85 | 1.03 | **1** |  | 0.7811/0 |
| AGP336a | *tcer-1* | 87/112 | 70.61 | 2.09 | **0** |  |  |
| AGP354 | *tcer-1;lipl-1* | 85/120 | 56.04 | 1.23 | **0.0004** | **0.0009** |  |
| AGP358 | *tcer-1;lipl-2* | 66/124 | 78.23 | 2.66 |  | **1** |  |
| AGP360a | *tcer-1; lipl-1 lipl-2* | 78/128 | 53.99 | 1.3 |  | **0.000029** | **0.000024/0** |

**Table S4:** Impact of *lipl-1* and *lipl-2* null mutants on lifespan on *E. coli* OP50.

| Strain | Background Genotype | Trial 1 |  |  | Bonferroni P-value |  |
| --- | --- | --- | --- | --- | --- | --- |
|  |  | n = obs/<br>total | Mean | SE ^ | P (vs N2) | P (vs *tcer-1*) |
| N2 | WT | 77/121 | 19.03 | 0.77 |  |  |
| AGP347 | *lipl-1* | 79/116 | 17.4 | 0.65 | 0.537 |  |
| AGP364a | *lipl-2* | 71/120 | 20.6 | 0.56 | 1 |  |
| AGP357 | *lipl-1 lipl-2* | 53/124 | 18.1 | 0.74 | 1 |  |
| AGP336a | *tcer-1* | 65/118 | 15.87 | 0.74 | 0.0665 |  |
| AGP354 | *tcer-1;lipl-1* | 72/115 | 19.36 | 0.79 |  | **0.0113** |
| AGP358 | *tcer-1;lipl-2* | 87/117 | 19.19 | 0.78 |  | **0.0297** |
| AGP360a | *tcer-1;lipl-1 lipl-2* | 82/117 | 15.75 | 0.5 |  | 1 |
| **Trial 2** |  |  |  |  |  |  |
| N2 | WT | 55/103 | 15.75 | 0.68 |  |  |
| AGP347 | *lipl-1* | 71/110 | 13.81 | 0.67 | 1 |  |
| AGP364a | *lipl-2* | 52/103 | 15.8 | 0.96 | 1 |  |
| AGP357 | *lipl-1 lipl-2* | 64/100 | 11.95 | 0.66 | **0.0004** |  |
| AGP336a | *tcer-1* | 39/103 | 12.08 | 0.76 | **0.0031** |  |
| AGP354 | *tcer-1;lipl-1* | 83/103 | 12.9 | 0.54 |  | 1 |
| AGP358 | *tcer-1;lipl-2* | 79/97 | 15.31 | 0.69 |  | **0.0369** |
| AGP360a | *tcer-1;lipl-1 lipl-2* | 80/104 | 13.78 | 0.72 |  | 0.5519 |
| **Trial 3** |  |  |  |  |  |  |
| N2 | WT | 94/121 | 14.96 | 0.34 |  |  |
| AGP347 | *lipl-1* | 80/106 | 15.79 | 0.57 | 0.2334 |  |
| AGP364a | *lipl-2* | 89/113 | 15.93 | 0.56 | 0.1141 |  |
| AGP357 | *lipl-1 lipl-2* | 101/114 | 13.84 | 0.48 | 1 |  |
| AGP336a | *tcer-1* | 92/120 | 14.23 | 0.48 | 1 |  |
| AGP354 | *tcer-1;lipl-1* | 77/120 | 15.36 | 0.51 |  | 0.9902 |
| AGP358 | *tcer-1;lipl-2* | 81/103 | 16.72 | 0.58 |  | **0.0133** |
| AGP360a | *tcer-1;lipl-1 lipl-2* | 86/114 | 13.84 | 0.35 |  | 1 |

**Table S5:** Impact of *lipl-1* and *lipl-2* knockouts on survival upon *S. aureus* infection.

| Genotype | # Animals | Mean (Hours) | S.E.M | P-value |  | Bonferroni P-value |  |
| --- | --- | --- | --- | --- | --- | --- | --- |
|  |  |  |  | vs. WT | vs. *tcer-1* | vs. WT | vs. *tcer-1* |
| Trial 1 |  |  |  |  |  |  |  |
| WT | 65 | 40.26 | 1.47 |  |  |  |  |
| *tcer-1* | 60 | 53.45 | 1.75 | <0.0001 |  | <0.0001 |  |
| *tcer-1; lipl-1* | 71 | 49.19 | 1.54 | <0.0001 | 0.95 | <0.0001 | 1 |
| *tcer-1;lipl-2* | 43 | 57.42 | 2.2 | <0.0001 | 0.03 | <0.0001 | 0.08 |
| Trial 2 |  |  |  |  |  |  |  |
| *WT* | 81 | 34.84 | 1.25 |  |  |  |  |
| *tcer-1* | 76 | 59.21 | 2.33 | <0.0001 |  | <0.0001 |  |
| *tcer-1; lipl-1* | 65 | 52.72 | 2.08 | <0.0001 | 0.16 | <0.0001 | 0.83 |
| *tcer-1;lipl-2* | 63 | 64.95 | 2.43 | <0.0001 | 0.28 | <0.0001 | 0.49 |
| Trial 3 |  |  |  |  |  |  |  |
| WT | 73 | 41.07 | 1.90 |  |  |  |  |
| *lipl-1* | 73 | 42.29 | 1.82 | 0.01 | <0.0001 | 0.02 | <0.0001 |
| *tcer-1* | 60 | 50.50 | 2.29 | <0.0001 |  | <0.0001 |  |
| *tcer-1; lipl-1 lipl-2* | 80 | 40.14 | 1.82 | 0.51 | <0.0001 | 1.00 | <0.0001 |
| Trial 4 |  |  |  |  |  |  |  |
| WT | 59 | **33.62** | 1.71 |  |  |  |  |
| *lipl-1* | 41 | **41.32** | 2.49 | 0.81 | 0.18 | 1 | 0.53 |
| *tcer-1* | 43 | **50.56** | 3.08 | 0.02 |  | 0.05 |  |
| *tcer-1; lipl-1 lipl-2* | 50 | **45.61** | 2.29 | 0.08 | <0.0001 | 0.24 | <0.0001 |
| Trial 5 |  |  |  |  |  |  |  |
| WT | 72 | 43.16 | 1.82 |  |  |  |  |
| *lipl-1* | 65 | 40.99 | 2.12 | 0.15 | 0.32 | 0.44 | 0.75 |
| *tcer-1* | 52 | 46.10 | 2.50 | 0.64 |  | 1.00 |  |
| *tcer-1; lipl1 lipl2* | 72 | 40.45 | 1.38 | 0.89 | 0.02 | 1.00 | 0.06 |
| Trial 6 |  |  |  |  |  |  |  |
| WT | 73 | 39.05 | 2.47 |  |  |  |  |
| *lipl-2* | 73 | 54.35 | 2.46 | <0.0001 |  | <0.0001 |  |
| *lipl-1 lipl-2* | 77 | 56.32 | 1.74 | <0.0001 |  | <0.0001 |  |
| Trial 7 |  |  |  |  |  |  |  |
| WT | 79 | 37.01 | 1.20 |  |  |  |  |
| *lipl-2* | 74 | 54.25 | 2.00 | <0.0001 |  |  |  |
| Trial 8 |  |  |  |  |  |  |  |
| WT | 81 | 58.45 | 1.38 |  |  |  |  |
| *lipl-2* | 58 | 64.06 | 1.11 | <0.0001 |  | <0.0001 |  |
| *lipl-1 lipl2* | 76 | 61.58 | 1.41 | 0.01 |  | 0.02 |  |
| Trial 9 |  |  |  |  |  |  |  |
| WT | 78 | 50.53 | 1.57 |  |  |  |  |
| *lipl-1 lipl2* | 75 | 59.75 | 1.48 | <0.0001 |  | <0.0001 |  |

**Table S6:**
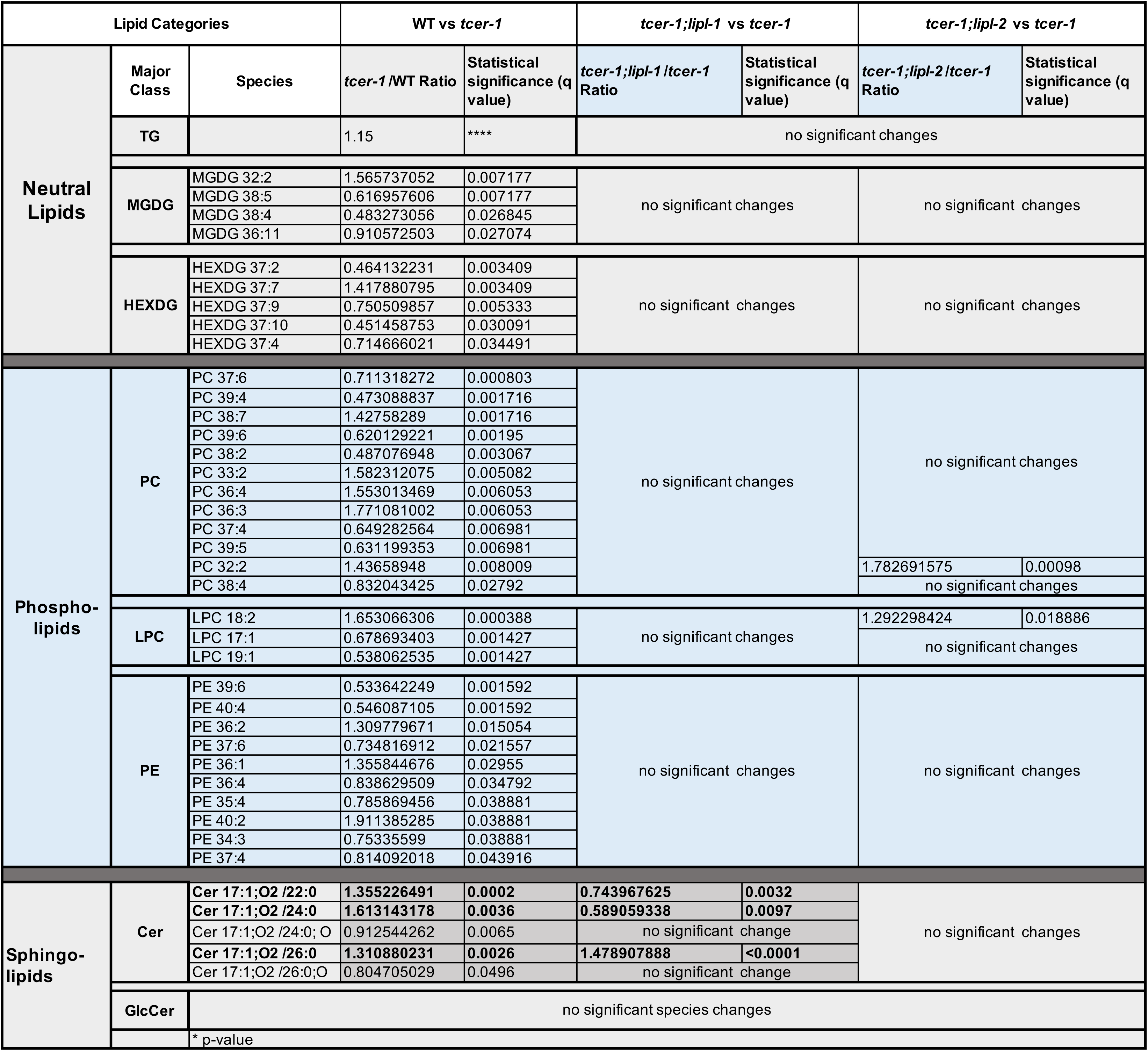
Lipid species altered in *tcer-1* mutants and impacts of *lipl-1* and *lipl-2* mutations on them.

**Table S7:**
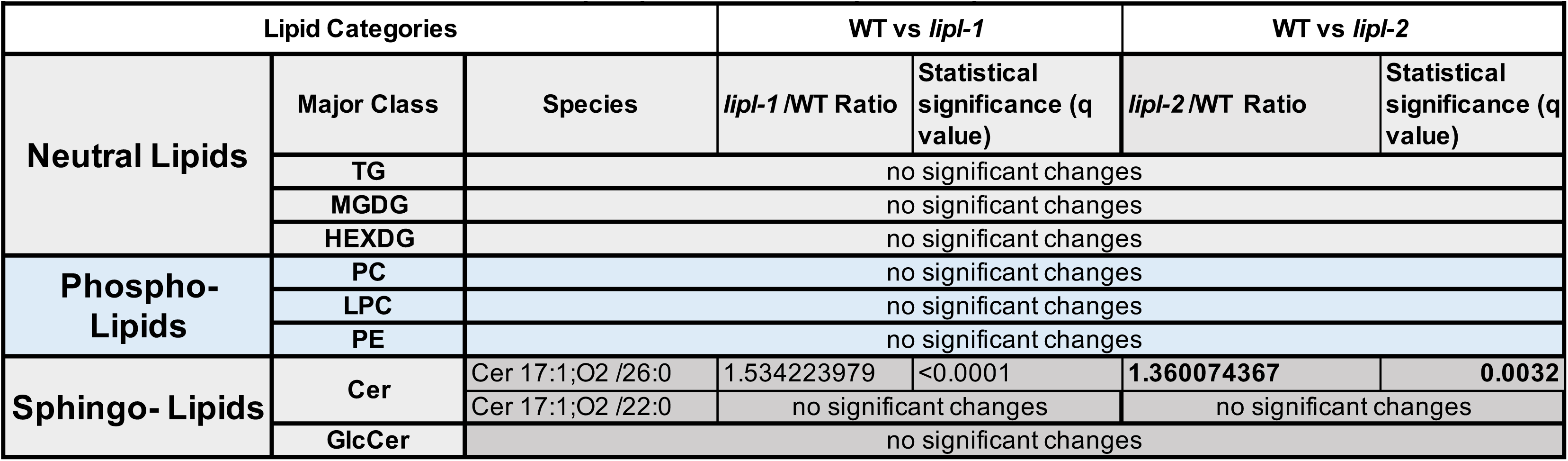
Lipid species altered in *lipl-1* and *lipl-2* mutants.

| Table S7: Lipid species altered in *lipl-1* and *lipl-2* mutants |  |  |  |  |  |  |
| --- | --- | --- | --- | --- | --- | --- |
| Lipid Categories |  |  | WT vs *lipl-1* |  | WT vs *lipl-2* |  |
| Neutral Lipids | Major Class | Species | *lipl-1* /WT Ratio | Statistical significance (q value) | *lipl-2* /WT Ratio | Statistical significance (q value) |
|  | TG | no significant changes |  |  |  |  |
|  | MGDG | no significant changes |  |  |  |  |
|  | HEXDG | no significant changes |  |  |  |  |
| Phospho-Lipids | PC | no significant changes |  |  |  |  |
|  | LPC | no significant changes |  |  |  |  |
|  | PE | no significant changes |  |  |  |  |
| Sphingo- Lipids | Cer | Cer 17:1;O2 /26:0 | 1.534223979 | <0.0001 | 1.360074367 | 0.0032 |
|  |  | Cer 17:1;O2 /22:0 | no significant changes |  | no significant changes |  |
|  | GlcCer | no significant changes |  |  |  |  |

**Table S8:** Survival of strains expressing human LAL (hLAL) in different genetic backgrounds.

| Strain | Genotype | Trial 1 |  |  | Bonferroni P-value |  |  |
| --- | --- | --- | --- | --- | --- | --- | --- |
|  |  | n = obs/<br>total | Mean | SE ^ | P (vs WT) | P (vs *tcer-1*) | P (vs. *tcer-1;lipl-1*) |
| WT* | WT | 54/90 | 35.06 | 1.91 |  | **0.0000034** | 1 |
| CF2166* | *tcer-1* | 63/90 | 54.8 | 3.25 | **0.0000034** |  | **0.000021** |
| AGP354* | *tcer-1;lipl-1* | 52/90 | 36.65 | 1.79 | 1 | **0.000021** |  |
| * | *tcer-1;lipl-1;hLAL (A)* | 73/90 | 55.77 | 2.53 | **0** | 1 | **0** |
|  | *tcer-1;lipl-1;hLAL (B)* | 70/90 | 40.14 | 1.68 | 0.0738 | **0.0002** | 0.5904 |
| Trial 2 |  |  |  |  |  |  |  |
| N2 | WT | 51/90 | 40.29 | 2.7 |  | **0** | **0.0000027** |
| CF2166 | *tcer-1* | 54/90 | 68.56 | 2.99 | **0** |  | **0.0126** |
| AGP354 | *tcer-1;lipl-1* | 42/90 | 55.31 | 4.57 | **0.0000027** | **0.0126** |  |
|  | *tcer-1;lipl-1;hLAL (A)* | 43/90 | 63.53 | 2.59 | **0** | 0.4523 | 0.0866 |
|  | *tcer-1;lipl-1;hLAL (B)* | 39/90 | 63.08 | 3.66 | **0** | 0.6723 | 0.1765 |

\* Shown in Figure 7A

**Table S9:**
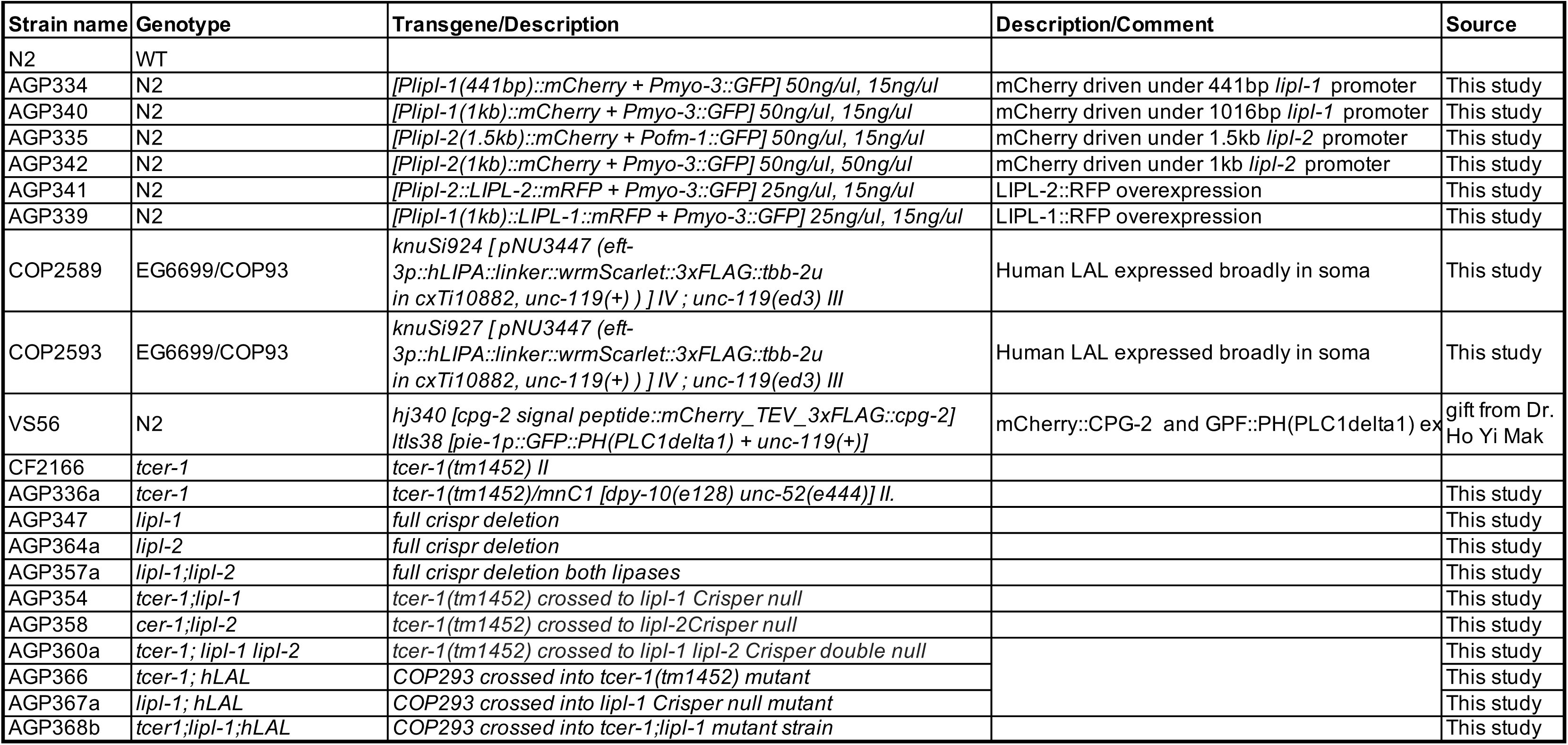
Strains used in this study.

**Table S10:**
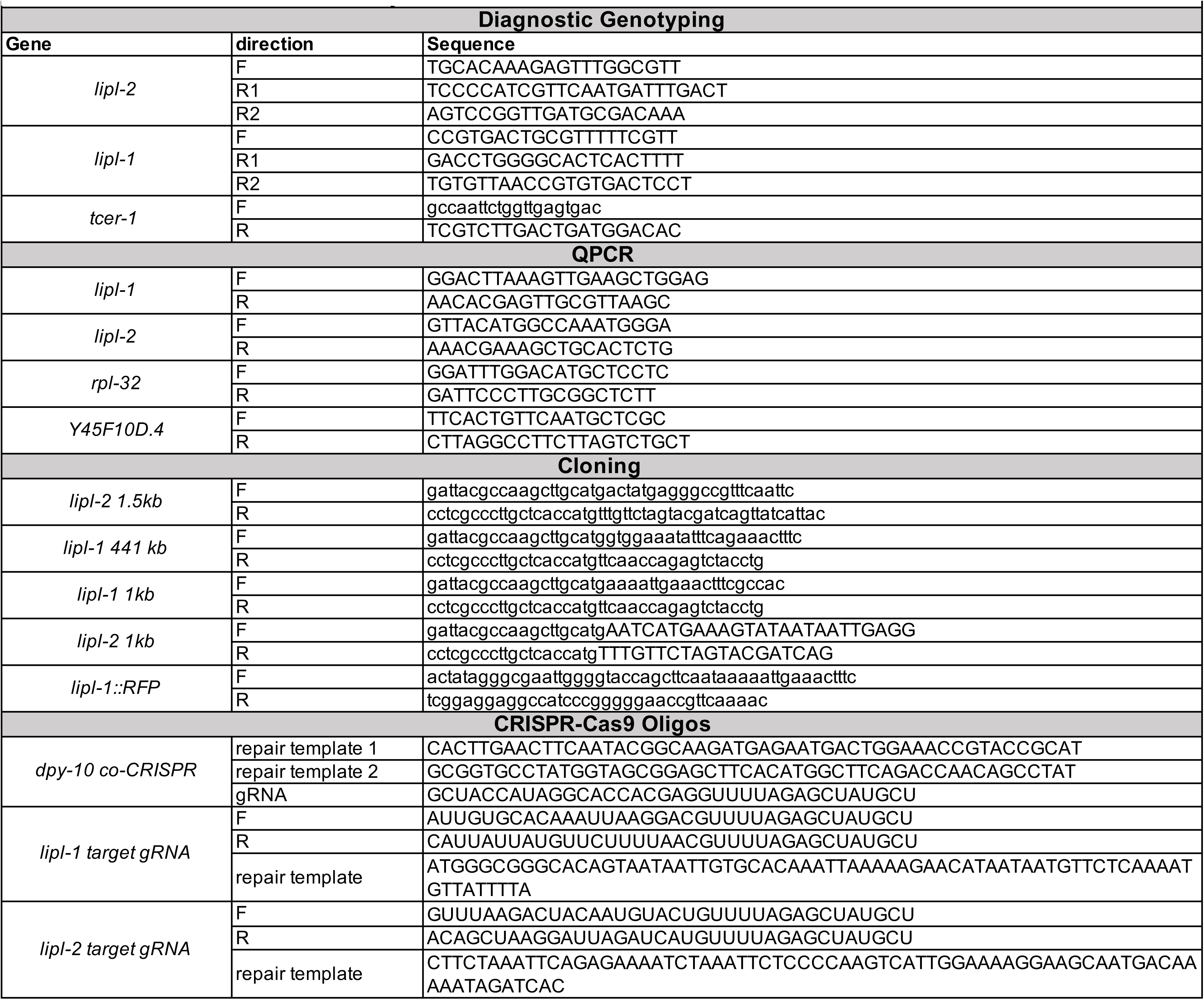
Primers used in this study.

